# tRNA isodecoder analysis using Nanopore ionic current signals and deep learning

**DOI:** 10.64898/2025.12.27.696718

**Authors:** Stuart Akeson, Pooria Daneshvar Kakhaki, Neda Ghohabi Esfahani, Julia L. Reinsch, Margaret L. Barry, Maddy Zamecnik, Talia Tzadikario, Robin L. Abu-Shumays, David M. Garcia, Kristin S. Koutmou, Miten Jain

## Abstract

tRNA are short non-coding RNA characterized by their distinct tertiary structure and abundant chemical modifications. Conventional analysis strategies do not fully characterize tRNA isodecoders. We demonstrate that this limitation can be resolved for tRNA using nanopore ionic current data. We developed tRNAZAP, a deep learning strategy that uses nanopore ionic current signal information to classify native tRNA strands at isodecoder-level resolution without relying on sequence information. Additionally, the ionic current level classification allows for pairwise alignment of read sequences to reference sequences, producing optimal tRNA alignments. We applied tRNAZAP to direct tRNA sequencing data from *Escherichia coli* and *Saccharomyces cerevisiae,* and recovered 2.6% and 13.1% more aligned reads than BWA-MEM, respectively. tRNAZAP resolved these reads at an isodecoder-level and with consistently higher alignment identity. tRNAZAP is a powerful complement to sequence-based profiling and can contribute towards resolving the isodecoder landscape in more complex organisms including humans.

## INTRODUCTION

tRNA are the most abundant RNA class in terms of molecular concentration. They are typically ∼76 nucleotides in length and include extensive chemical modifications^1,2^. tRNA research is predominantly focused on their abundance, sequence, aminoacylation status, and chemical modifications. Techniques for studying these features include Northern blots^3–5^, liquid chromatography paired with tandem mass spectrometry (LC-MS)^2,6–8^, and various short-read sequencing technologies that require reverse transcription and PCR for generating cDNA copies^8,9^. Recently, Nanopore direct RNA sequencing has emerged as a powerful technique for studying tRNA, especially because it can provide direct measurement of RNA modifications at a single molecule level^6,10–12^.

Due to the structured and highly modified nature of tRNA, the use of complementary methods can help resolve aspects of tRNA biology^6,8^. For example, we previously combined Nanopore direct RNA sequencing, LC-MS, and genetic controls to study the coordination among T-loop modifications in *Saccharomyces cerevisiae* (*S. cerevisiae*)^6^. These approaches can help advance tRNA research including modifications that have been linked to cancer, metabolic disorders, and neurodegenerative disease^1^.

tRNA are typically categorized into three categories: i) Isotypes – different tRNA that carry the same amino acid; ii) Isoacceptors – different tRNA that carry the same amino acid but have distinct anticodon sequences; and iii) Isodecoders – tRNA that carry the same amino acid and have the same anticodon sequence but have small sequence differences in the tRNA “body” (any nucleotide other than the anticodon)^13^. Isodecoders are the most challenging to distinguish because they can differ by as little as a single nucleotide^10^. As a result, the majority of tRNA sequence analysis relies on isoacceptor-level discrimination^6,12^. Merging isodecoders under a single isoacceptor label simplifies biological complexity. Disambiguation of these isodecoders in sequence space remains an ongoing challenge^14^.

Prior work demonstrated that Nanopore direct RNA sequencing (DRS) can provide full-length readouts of native tRNA from both prokaryotes and eukaryotes^6,12,15–17^. During Nanopore sequencing, a tRNA passes through the protein pore, across which voltage has been applied. The specific order of nucleotides produces a series of unique ionic current signatures^18^. Decoding these ionic signatures back into a sequence of nucleotides is called basecalling. The process of basecalling tRNA is challenging because of the dense modifications. Basecalling accuracy can be measured using alignment identity, a process that compares the basecalled sequence to a known reference sequence. Nanopore based tRNA alignment identity ranged between 82.35% to 89.47% for *S. cerevisiae* and *E. coli* respectively. This is lower than the current median alignment identity of 98.7% for human mRNA^19–21^. This impact on accuracy is, in part, due to the presence of complex and dense chemical modifications in tRNA. It has been documented that RNA modifications can cause perturbations in Nanopore ionic current^18,22–24^. Furthermore, these perturbations affect the accuracy of the basecalling step. This observation has been previously exploited for RNA modification analysis^6,15,25–31^.

Nanopore tRNA sequence analysis typically utilizes BWA-MEM for alignment. BWA-MEM allows for rapid alignment of short nucleic acid sequences across a large set of reference sequences. But BWA-MEM is a heuristic aligner, meaning that it does not always produce the optimal alignment. This, coupled with 80-90% sequence alignment identity, makes differentiation of isodecoders at the single nucleotide level incomplete. To address this, researchers, including ourselves, have previously used limited-scope references that collapse all isodecoders under a single isoacceptor reference sequence^6,12,15^. A recent strategy to disambiguate isodecoders used a hierarchical alignment approach. This strategy attributes reads to increasingly refined classifications of tRNA moving from a given isotype to the underlying isodecoder depending on their alignment scores^14^. This method of limiting classification is principled, but purely a sequence-level approach and achieved isodecoder-level assignment for ∼30% of human tRNA mapped reads. Improving isodecoder resolution will allow for increased understanding of tRNA biology.

Recent efforts have demonstrated that ionic current analysis can be used for tRNA specific applications. These include: i) measuring the aminoacylation status of tRNA^16^; ii) applying ionic current-based modification models trained for mRNA for “off-label” modification predictions in tRNA^11^; and iii) ionic current classification for demultiplexing barcoded tRNA^32^. Despite this progress, the isodecoder disambiguation challenge continues^12,15^. In this work, we demonstrate a novel ionic current-based strategy (tRNAZAP) that achieves isodecoder-level resolution in *E. coli* and *S. cerevisiae* tRNA. tRNAZAP uses ionic current signal information to provide isodecoder-level classification, tRNA-specific ionic current segmentation, and a pairwise alignment step that uses both sequence and ionic current information for optimal tRNA sequence alignment. We document significant improvements in alignment of tRNA reads compared to previous methods for *E. coli* and *S. cerevisiae*. We also demonstrate discrimination among closely-related isodecoders using ionic current analysis. tRNAZAP outputs tRNA sequence alignments in BAM format, making them suitable for conventional downstream genomic analyses. tRNAZAP allows for tRNA analysis predominantly using ionic current rather than sequence information. This is in line with the broader field of RNA modification analysis that has also started employing ionic current-based modification calling using deep learning^19,33^.

## RESULTS

### In vitro transcription of Escherichia coli tRNA

We performed *in vitro* transcription for the complete set of non-redundant *Escherichia coli* (*E. coli)* tRNA isodecoders predicted by gtRNAdb^34^ (**Supplementary Table 1**). To facilitate ionic current learning we augmented our previous adapter design^6,15^ to include a longer 3′ component (**Supplementary Table 2)**. We sequenced the in vitro transcribed (IVT) tRNA using Oxford Nanopore Technologies (ONT) RNA004 platform on a PromethION flow cell. This sequencing experiment produced 18,985,579 reads. We aligned the reads using BWA-MEM (-x ont2d -w 13 -K 6). Of these, 15,310,260 (80.64%) reads were primary alignments (mapping quality over 0). Those reads had a median alignment identity of 97.26% and a median aligned read length of 140 bases (**Supplementary Figure 1**). The high alignment identity of the IVT tRNA supports the conclusion that tRNA modifications are the primary drivers for lower tRNA alignment identities. We documented a median of 108,422 reads amongst the 51 unique *E. coli* tRNA isodecoders. *E. coli* Gln-TTG-1-1 (1,086) and *E. coli* Leu-TAA-1-1 (256) had fewer than 2,000 reads and were resequenced separately. In total, we documented a minimum of 39,067 primary reads with a mapping quality greater than 0 across all 51 isodecoders (**Supplementary Table 3**).

### Ionic current model training using IVT *E. coli* tRNA data

We used the IVT tRNA DRS data to train tRNAZAP (a transformer-based model). Transformers are a class of neural networks that learn complex patterns in sequential data. They achieve increased accuracy by focusing on the most informative parts of the input, including long-range signal interactions^35^. The training set for the tRNAZAP model was composed of independent reads equally balanced across all isodecoder classes (see Methods). The model was designed to produce two outputs simultaneously:

i) classification of ionic current traces into tRNA isodecoders; and
ii) segmentation of ionic current traces into the components of an adapted tRNA including the ONT sequencing adapter, tRNA splint 1 (ligated to tRNA 3′-end), tRNA isodecoder, and tRNA splint 2 (ligated to tRNA 5′-end).

Figure 1 illustrates an example output of tRNAZAP from the raw ionic current input. The signal segmentation provided by the tRNAZAP allows for a targeted input of sequence into the downstream pairwise sequence alignment algorithm. The Dorado moves table is used for mapping signal to sequence at a per-read level.

**Figure 1.**
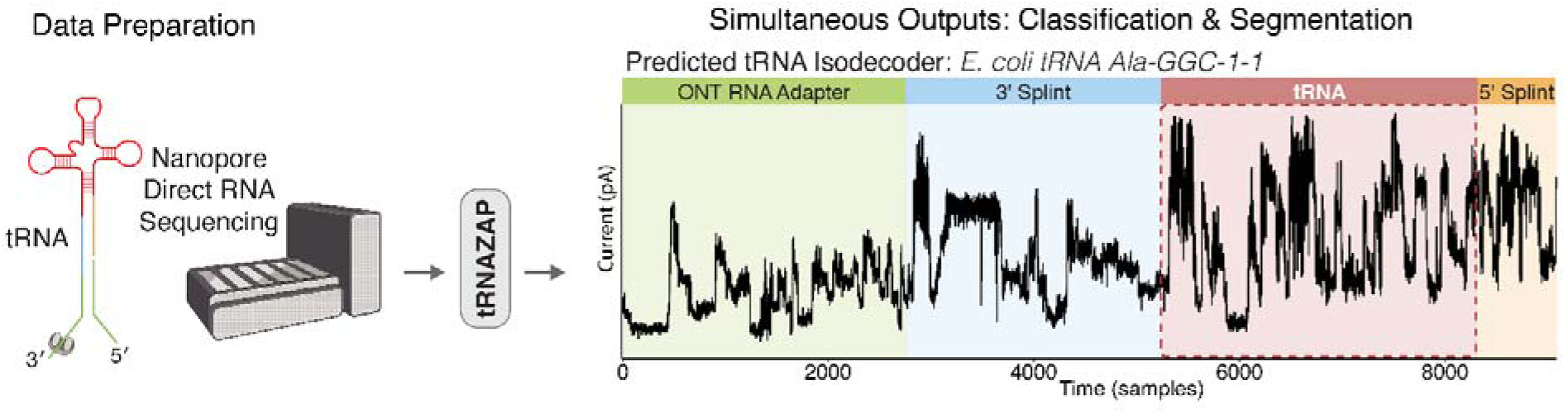
An overview of the tRNAZAP transformer-based end-to-end pipeline for tRNA analysis. The model takes an input ionic current signal (in picoamperes), simultaneously classifies the corresponding tRNA isodecoder and segments the signal into biologically relevant regions.

We first tested tRNAZAP on *E. coli* IVT tRNA data **(Supplementary Figure 2)**. We assessed tRNAZAP performance for both tRNA isodecoder classification and tRNA region segmentation. We used BWA-MEM alignment-based labels as the reference for this assessment (see Methods). For tRNA isodecoder classification, the model’s unweighted average accuracy across isodecoders was 99.16% (**Supplementary Figure 2A**). The overlap between BWA labeled signal and tRNAZAP segmented signal, calculated as the intersection over union (IoU), was 0.96 (**Supplementary Figure 2B**).

To assess if we could document ionic current differences between isodecoders, we used the dorado move-table to compare the ionic current alignments for two IVT tRNA isodecoders from the training data (Leu-CAG-1-1 and tRNA Leu-CAG-2-1). This isodecoder pair represents a challenging disambiguation due to the single nucleotide variance between the pair at reference position 47 (V13 using Sprinzl coordinates^36,37^). While the ionic current data suggest two distinct populations (Figure 2A), the single position separation is subtle. We then applied Uniform manifold approximation and projection (UMAP), a method for visualizing data in high dimensions, to test if the isodecoder specific reads were clustering separately (Figure 2B). It should be noted that UMAP prioritizes keeping closely related points close together rather than preserving global structure. The combined effect of ionic current shift in the region of the single nucleotide variant produced two clear populations in the UMAP visualization. This suggests that it is possible to separate isodecoders that differ by a single nucleotide for modification-free in vitro transcribed tRNA.

**Figure 2.**
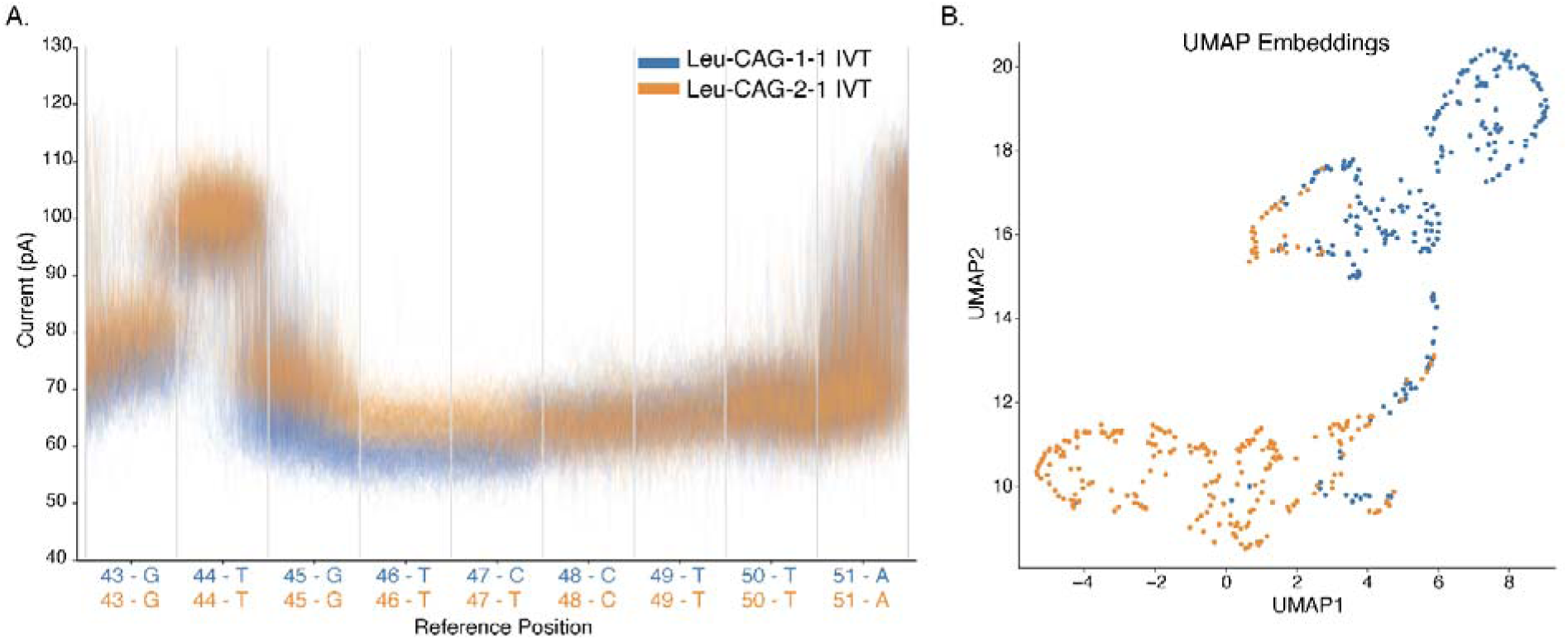
Ionic current profiles for ***E. coli*** IVT Leu-CAG isodecoders. **A.** Raw ionic current profiles for ***E. coli*** Leu-CAG isodecoders. The x-axis shows a 9 nucleotide length window centered on position 47 (V13 in Sprinzl coordinates) of tRNA Leu. Each segment of the x-axis corresponds to one nucleotide. The y-axis represents the raw ionic current signal in picoamperes. The colors correspond to the isodecoders, blue for Leu-CAG-1-1 and orange for Leu-CAG-2-1. The signal to sequence mapping was produced using the dorado moves table and was anchored to the reference sequence using BWA-MEM alignment. **B.** For UMAP, we took the median pA and standard deviation of each position from 45 to 47 for every read and plotted the 2D representation of those ionic current statistics. The coloring for the UMAP matches that of the raw ionic current plots.

### Model tuning using biological *E. coli* tRNA data

tRNAZAP performed well on IVT tRNA test data. However, because the IVT tRNA strands lacked chemical modifications, they did not fully represent the complexity of biological tRNA strands. We reasoned that biological tRNA data could be used to refine the ionic current model. This technique of refining a pre-trained model on new data, known as fine-tuning, is a standard practice in deep learning^38–40^. To that end, we sequenced *E. coli* total tRNA and assigned isodecoder-level tRNA labels using BWA-MEM under stringent alignment criteria (see Methods). We fine-tuned the model using these labeled biological tRNA data. The refined model showed higher concordance with BWA-MEM classification of labels for biological data (**Supplementary Figure 3, Supplementary Table 4**)

### An ionic current guided pairwise sequence alignment strategy

tRNAZAP performs isodecoder level classification for every strand based exclusively on ionic current signal. This removes the inability to distinguish closely related references, an issue common with conventional tRNA sequence alignment strategies^14^. With only a single reference option to consider, we implemented a computationally tractable pairwise alignment strategy (See Methods). This strategy generates an optimal alignment between the nanopore read sequence and the corresponding isodecoder reference sequence^41^. Pairwise alignment algorithms typically are computationally expensive^41^. This is mitigated by tRNAZAP-based isodecoder prediction which provides a single reference option for each read sequence. tRNAZAP further limits the computational burden of pairwise alignment by identifying the ionic current signal region associated with tRNA and only aligning the corresponding portion of sequence to the reference. To extract the tRNA relevant sequence we used the tRNAZAP segmentation coordinates coupled with the dorado move-table, an approximate signal to sequence mapping produced by the dorado basecaller.

For biological tRNA data, we used a sequence identity cut-off threshold of 75%, with a minimum of 25 matching bases in the tRNA region of the reference sequence. The length of 25 was selected to accommodate the potential for tRNA fragments split at the anticodon loop^42^, which can account for the characteristic loss of the ∼10-12 5′-terminal nucleotides in nanopore DRS^43^.

### Assessing tRNAZAP performance for biological *E. coli* tRNA

To test the tRNAZAP *E. coli* model we sequenced total tRNA from *E. coli* strain MRE600. Nanopore DRS produced a total of 51,034,636 reads. We filtered out reads with the ‘pi’ tag, which denotes read splitting: a step during basecalling by which Dorado tries to retrospectively fix when the ionic current signals for two consecutive strands are incorrectly recorded as one strand. This filtering reduced the number of reads to 50,753,889. tRNAZAP aligned 19,079,732 (37.59%) of these reads. This was 510,879 more reads than BWA-MEM (Figure 3A, D^2^=24,124.58, p<1×10^-308^). Additionally, tRNAZAP aligned reads with 89.37% alignment identity, 0.37% higher than BWA-MEM (Figure 3B, paired Wilcoxon signed-rank test; W = 9.05×10^12^, p<1×10^-308^). This means that using tRNAZAP in place of BWA-MEM is producing more reads with higher alignment identity per sequencing experiment.

**Figure 3.**
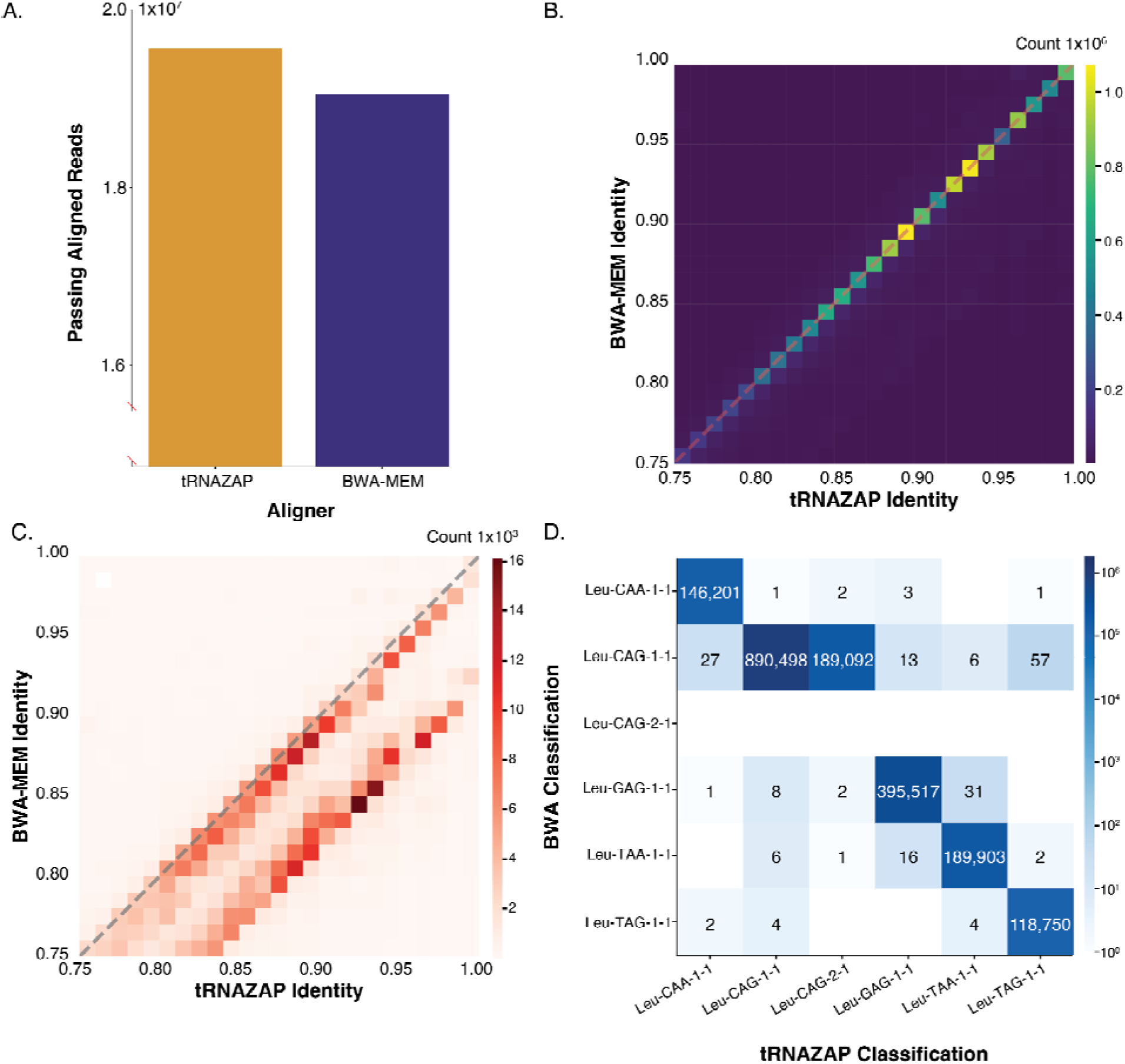
tRNAZAP performance for *E. coli* biological tRNA compared with BWA-MEM **A.** Count of reads (y-axis) that achieved a minimum of 25 matching bases in the tRNA region and a minimum of 75% alignment identity for tRNAZAP and BWA-MEM (x-axis). **B.** Per read alignment identity for the tRNA portion of the reference compared at a paired read level between tRNAZAP and BWA-MEM. Colors represent the raw count of reads per bin. **C.** Per read alignment identity compared between tRNAZAP and BWA-MEM for reads where the read aligned to different primary tRNA. Colors represent the raw number of reads per bin. **D.** A confusion matrix showing the per isodecoder counts of reads for *E. coli* tRNA isotype Leucine and the 6 corresponding isodecoders spread across 5 isotypes. The only isodecoder pair is Leu-CAG-1-1 and Leu-CAG-2-1. This is not the complete set of possible alignments, but rather a subset for demonstrative purposes. The BWA-MEM reference does not include Leu-CAG-2-1 explaining the complete lack of reads present in this row. The x-axis shows the reference that tRNAZAP aligned a given read to, while the y-axis shows the reference that BWA-MEM aligned a given read to. Both of these aligners are computational techniques applied to the same set of reads, so every read will have a tRNAZAP classification and a BWA-MEM classification. The combination of these references determines which box receives the count for a read. The color of each box as well as the text in each box show the total number of reads that have a given pairing of selected references. Boxes with no coloration nor text have no reads aligning to them.

BWA-MEM and tRNAZAP disagreed on the assignment of reference sequences for 4.12% of the reads (**Supplementary Table 5, Supplementary Figure 4)**. tRNAZAP achieved an alignment identity of 88.45% for these reads, similar to the earlier alignment identity. In contrast, BWA-MEM achieved 84.48% alignment identity for this set (-3.97%) (Figure 3C, paired Wilcoxon signed-rank test; W = 2.43×10^11^, p<1×10^-308^). Figure 3D illustrates the counts of reads that were aligned to the tRNA isotype Leucine for both BWA-MEM and tRNAZAP. Reads that were classified differently by tRNAZAP and BWA-MEM were counted in the boxes off of the diagonal (top left to bottom right). Of these boxes 99.90% were attributed to Leu-CAG-2-1, an isodecoder that had to be removed from the BWA-MEM reference set to prevent multimapping. It is logical that given a set of references that include isodecoders, and the ability to distinguish between those isodecoders, an aligner would achieve higher alignment identity. This means that tRNAZAPs ability to distinguish isodecoders helped align more reads at a higher alignment identity than BWA-MEM.

We anticipated that properly separated biological isodecoder pairs would be associated with distinct ionic current profiles as was observed for IVT tRNA isodecoders. To test this, we plotted the raw ionic current signals (Figure 4A) for the same set of isodecoders that we had examined in IVT *E. coli* tRNA data. We noted that the ionic current profile was not only distinct between the biological isodecoders, but it was also similar in appearance to the IVT traces (Figure 2A) Extending the ionic current visualization with a UMAP projection accentuated the two distinct populations visible in the raw signal (Figure 4B). This evidence suggests that we are capturing two distinct tRNA populations across the two isodecoders when using ionic current signal information. When we performed a UMAP projection on both biological and IVT conditions simultaneously, we observed two populations corresponding to the two isodecoders **(Supplementary Figure 5)**. Each of the isodecoder types was populated by both biological and IVT tRNA. This provides evidence that the ionic current pattern, and by extension tRNAZAP’s learning, are providing isodecoder resolution.

**Figure 4.**
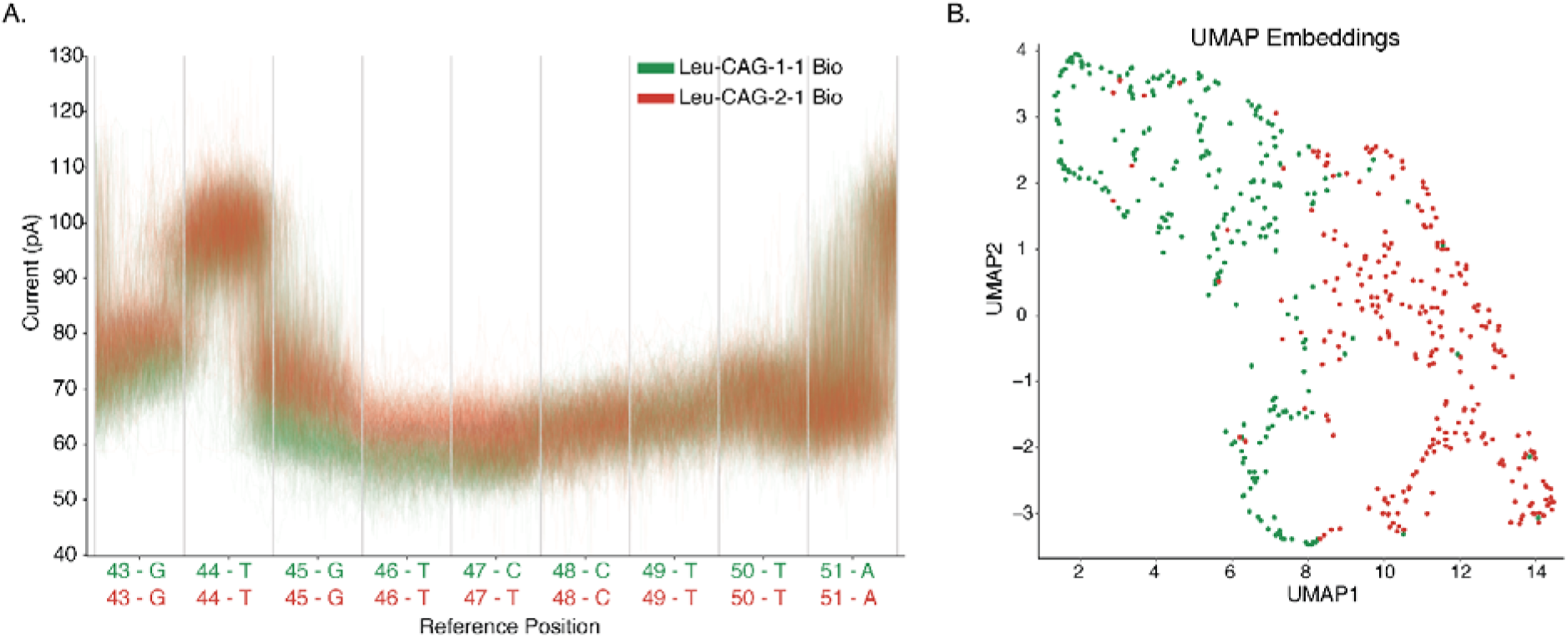
Ionic current profiles for ***E. coli*** biological Leu-CAG isodecoders. **A.** Raw ionic current profiles for ***E. coli*** Leu-CAG isodecoders. The x-axis and y-axis are the same for this figure as **Figure 2A** to allow for comparison. **B.** For UMAP we took the median pA and standard deviation of each position from 45 to 47 for every read and plotted the 2D representation of those ionic current statistics. The coloring for the UMAP matches that of the raw ionic current plots. It should be noted that UMAP coordinates are not comparable between figures.

### Assessing tRNAZAP on a biological *E. coli* tRNA sample enriched for tRNA^Phe^

It is possible for deep learning models to overfit to training data. We tested if tRNAZAP could capture isodecoder populations that differed from wild type, such as a purified subset of tRNA. There are several methods for enriching and analyzing a specific set of tRNA or even individual tRNA populations^44,45^. Assessing these enrichments would be critical for a broadly applicable Nanopore-based tRNA analysis strategy. We tested tRNAZAP for a non-wild-type condition that used HPLC purification to enrich *E. coli* tRNA Phe-GAA-1-1. We adapted and sequenced this fraction using the same protocol as the wild-type experiment. We documented a Fold Change (FC) of 17.01 transcripts per million (TPM) for Phe-GAA-1-1 between the total tRNA condition and the enrichment condition (Figure 5, **Supplementary Table 6**). When we examined the normalized read counts between the two conditions, we observed the expected enrichment of Phe-GAA-1-1, but also increased levels of Glu-TTC-1-1 (2.59 FC of TPMs), Ser-GCT-1-1 (4.78 FC of TPMs), and Ser-GGA-1-1 (3.47 FC of TPMs). The co-enrichment was also shown in BWA-MEM alignments (**Supplementary Figure 6**), suggesting this is an artifact of the purification, not specific to tRNAZAP. This demonstrated that tRNAZAP was not overfitted during training and could be used for targeted tRNA population analyses.

**Figure 5.**
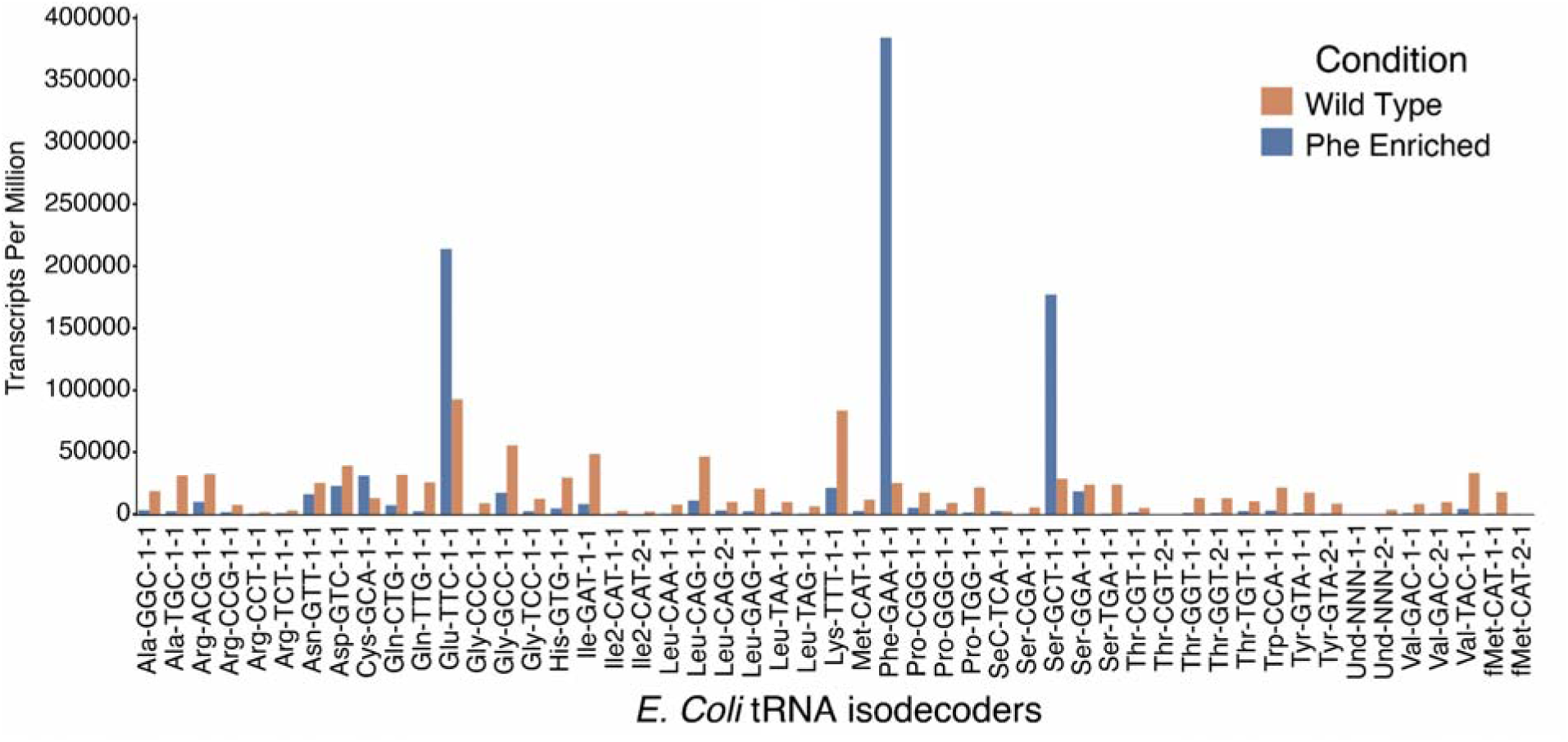
tRNAZAP performance on wild-type and tRNA^Phe^ enriched *E. coli* tRNA samples.The y-axis indicates Transcripts per million (TPM) while the x-axis shows the 51 *E. coli* isodecoders. Blue bars show the normalized tRNAZAP counts for the enriched tRNA^Phe^ condition while orange bars show the counts for the non-enriched wild type condition.

### Training tRNAZAP for a model eukaryote

Eukaryotic tRNAs contain more chemical modifications than *E. coli,* as profiled in previous studies^46^. To test tRNAZAP for eukaryotic tRNA, we trained a separate model for budding yeast (*S. cerevisiae*) tRNA. *S. cerevisiae* has a higher isodecoder-level complexity than *E. coli*. For example, the *E. coli* tRNA reference set includes 6 isoacceptors, each with two isodecoders. In contrast, the *S. cerevisiae* reference set includes 11 isoacceptors that correspond to a total of 23 isodecoders. Notably, the *S. cerevisiae* isoacceptor Gln-TTG has three isodecoders.

We generated an IVT tRNA-based model for *S. cerevisiae* using the same strategy that we had employed for *E. coli*. This model achieved similar performance (98.83% macro-average-accuracy ; 0.98 median IoU) to the *E. coli* IVT tRNA model. We fine tuned the model using data from three technical replicates of yeast biological tRNA sequencing data. During fine tuning we noted the increased complexity in modifications impacted basecalling quality and decreased alignment identity when using BWA-MEM (**Supplementary Table 7**).

We tested the fine-tuned *S. cerevisiae* model on data from an independent sequencing experiment. We used the same filtering thresholds for our comparison of BWA-MEM and tRNAZAP for our *S. cerevisiae* model as we did for the *E. coli* comparison. The *S. cerevisiae* sequencing run produced a total of 34,133,760 reads. Among those, we used 34,012,875 reads after removing strands that underwent read splitting. tRNAZAP aligned 10,173,310 (30.19%) of these reads under the same filtering thresholds. This was 1,179,020 reads more than BWA-MEM, a statistically significant 13.11% increase (Figure 6A, χ^2^ = 119551.89, p<1×10^-308^).

**Figure 6.**
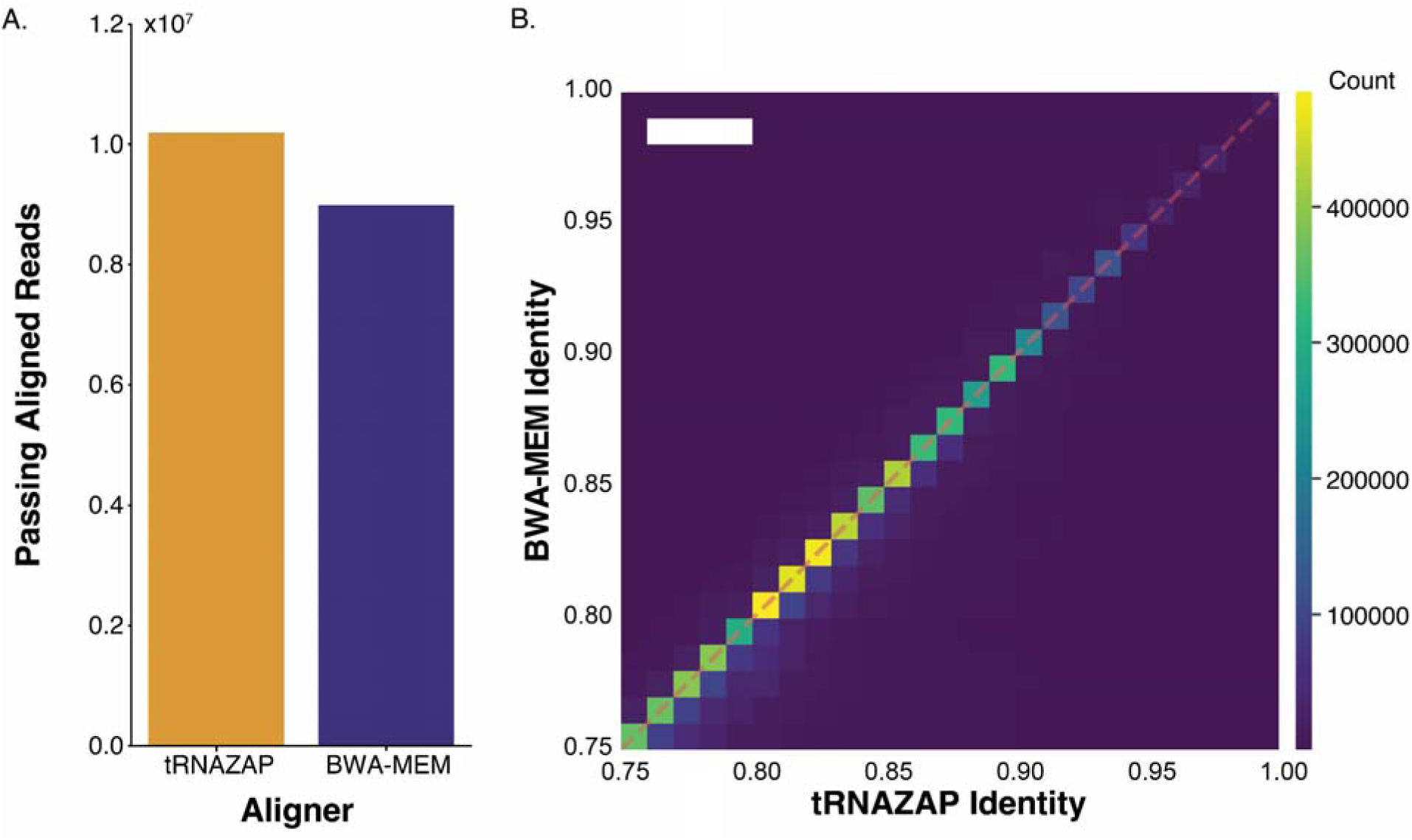
tRNAZAP performance for *S. cerevisiae* tRNA nanopore sequencing data. **A.** Count of reads (y-axis) that achieved a minimum of 25 matching bases in the tRNA region and a minimum of 75% alignment identity for tRNAZAP and BWA-MEM (x-axis) for *S. cerevisiae* biological tRNAs. **B.** Per read alignment identity for the tRNA portion of the reference compared at a paired read level between tRNAZAP and BWA-MEM. The x-axis shows the tRNAZAP alignment identity for a given read while the y-axis shows the corresponding BWA-MEM alignment identity for the same read. Colors in each box represent the raw count of reads per bin.

For reads aligned by both aligners, tRNAZAP achieved an alignment identity of 83.44% while BWA-MEM was 0.49% lower. This again demonstrated a statistically significant advantage for tRNAZAP alignment identity compared to BWA-MEM (Wilcoxon ranked-sign W=4.66×10^13^, p<1×10^-308^). For reads where both aligners had a successful alignment we documented a mean alignment identity of 82.95% and 83.44% for BWA-MEM and tRNAZAP respectively. We performed a paired Wilcoxon signed-rank test that confirmed that the observed 0.49% median identity increase for the shared aligned reads was significant (W=6.31×10^12^, p<1×10^-308^, Figure 6B). While the shifts we have observed in both *E. coli* and *S. cerevisiae* are proportionally small, they reinforce the observation that across species tRNAZAP is providing consistently higher alignment identity.

### tRNAZAP resolves isodecoders in *E. coli* and *S. cerevisiae* data

Isodecoder-level resolution of aligned tRNA read sequences has been challenging for Nanopore DRS^12,15,34^. This is due to the limited number of sequence differences between isodecoders and impact of modifications on sequencing accuracy. Prior nanopore tRNA analysis approaches, including our work, focused on isoacceptor-level analyses using curated reference sequences^6,10,12,15^. This loss of information oversimplifies tRNA biology and can obscure disease relevant discoveries. For instance, it has been shown that a single base mutation in the T-loop of mouse tRNA-Arg-UCU isodecoder encoded by tRNA gene *n-Tr-20* can cause fatal neurodegeneration^47^. Analysis strategies that resolve isodecoders at a single-molecule level will advance tRNA research.

tRNAZAP fills this gap by classifying reads at an isodecoder level. However, it is important to point out that our source of isodecoder level fine-tuning labels was based on BWA-MEM alignments. To ensure our training sets were as representative of specific isodecoders as possible, we only included reads whose sequence matched isodecoder specific positions with 100% agreement (See Methods). This technique has some risk of introducing a BWA-MEM based alignment bias into model training. An alternative approach would require the purification and direct RNA sequencing of individual isodecoders^45^. This is experimentally challenging and, due to the lack of accessible and published techniques, beyond the scope of the current study. Additionally, it is an open question if this would provide a purer training set than the lossy sequence level assignment we used.

This led to the question of how well tRNAZAP assigned isodecoder classes in the absence of ground truth labels. To assess this, we examined two independent metrics: alignment identity statistics and isodecoder specific ionic current profiles. Our rationale is that correct isodecoder level assignment by the model should produce distinguishable ionic current patterns and increase alignment identity due to pairing sequences with the proper reference.

For alignment identity statistics we earlier demonstrated that *E. coli* had significantly higher alignment identity for the reads where tRNAZAP and BWA-MEM disagreed (Figure 3C). This supports the successful disambiguation of *E. coli* isodecoders. We applied the same logic to *S. cerevisiae* isodecoders. We identified the set of reads where tRNAZAP and BWA-MEM disagreed for *S. cerevisiae,* including reads that were assigned to an isodecoder BWA-MEM did not have in the reference (**Supplementary Table 8, Supplementary Figure 7**). In total, this set included 446,378 reads, of which 348,173 were assigned to an isoacceptor by BWA-MEM but a different isodecoder under the same isoacceptor by tRNAZAP. These 348,173 reads, if accurate, represent the isodecoder resolution capability of tRNAZAP. However, analyzing the alignment identity of these reads alone would favor tRNAZAP by excluding reads the model may have misclassified more broadly. To provide a fair assessment, we analyzed the full set of 446,378 alternatively classified reads tRNAZAP. These reads had a mean alignment identity of 82.40% compared to BWA-MEM’s 80.92% identity. This means tRNAZAP had a 1.48% higher mean identity, which was statistically significant (W=7.25×10^10^, p<1×10^-308^, Figure 7A). Figure 7B shows a confusion matrix for *S. cerevisiae* tRNA isotype Arg aligned using the two different aligners. The off-diagonal reads, which represent disagreement between tRNAZAP and BWA-MEM. These reads can be predominantly attributed to the isodecoder level resolution of tRNAZAP distinguishing between Arg-ACG-1-1 and Arg-ACG-2-1. In total, the 348,173 isodecoder level reads in disagreement with BWA-MEM accounted for 78.0% of off-diagonal class assignments.

**Figure 7.**
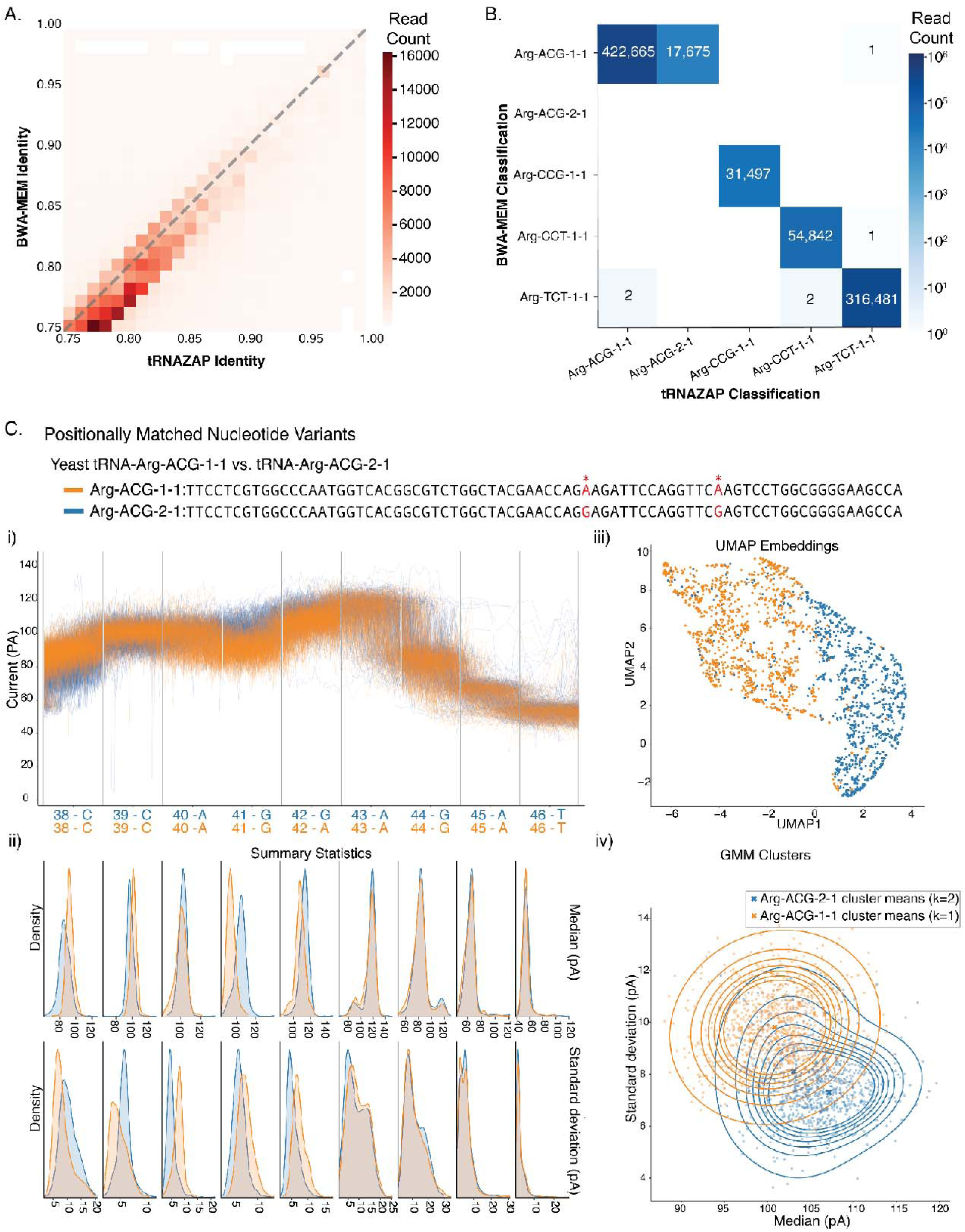
tRNAZAP Isodecoder-level resolution examination through alignment identity and ionic current profiles. **A.** 2D histogram depicting the alignment identity from tRNAZAP and BWA-MEM for the set of reads where the two aligners disagree on the best tRNA reference. The x-axis shows the tRNAZAP identity for a given read while the y-axis shows the corresponding BWA-MEM alignment identity. The reads for this heatmap are either i.) unmapped by one aligner, ii.) different isodecoders of the same isoacceptor, iii.) or are different isoacceptors or isotypes completely. Because the aligners are purely computational each read is aligned with both BWA-MEM and tRNAZAP allowing for paired comparison. **B.** Confusion matrix showing the per isodecoder counts of reads for *S. cerevisiae* tRNA isotype Arginine and the 5 corresponding isodecoders spread across 4 isotypes. The only isodecoder pair is Arg-ACG-1-1 and Arg-ACG-2-1. The BWA-MEM reference does not include Arg-ACG-2-1. The color of each box as well as the text in each box show the total number of reads that have a given pairing of selected references. Boxes with no coloration nor text have no reads aligning to them. **C.** Ionic current comparison between isodecoder variants tRNA-Arg-ACG-1-1 and tRNA-Arg-ACG-2-1. The two isodecoders differ at positions 42 and 55 (red asterisks). (**i**) Raw ionic current traces overlaid across nine positionally matched nucleotide positions centered on the variable base (G,A), colored by isodecoder. (**ii**) Kernel density estimates of per-position median (top) and standard deviation (bottom) of ionic current (pA) for each isodecoder. (**iii**) UMAP projection of multi-position ionic current features, revealing population structure between isodecoders. (**iv**) Two-dimensional Gaussian mixture models (GMMs) fitted to median and standard deviation features, with cluster means shown for each isodecoder (JS divergence = 0.48).

In addition to alignment statistics, we examined whether single-nucleotide differences between isodecoders produce distinguishable ionic current signatures. Using tRNA-Arg-ACG-1-1 and tRNA-Arg-ACG-2-1 as an example pair (differing at only two positions), we visualized and modeled their ionic current profiles across a 9-mer window centered at one of the varying positions. Overlaying the raw ionic current traces revealed distinct signal patterns between the two isodecoders (Figure 7C**.i**), most notably prior to the varying base at position 42 (A, G). To assess statistical significance, we used kernel density functions for the median and standard deviation of ionic current at each position for the reads across both populations. These kernel density estimates captured the distribution differences between the two isodecoders at each position, reinforcing the patterns observed in the ionic current traces (Figure 7C**.ii**). We then applied UMAP dimensionality reduction to project the multi-position feature space into two dimensions, where the two isodecoder populations formed distinct clusters (Figure 7C**.iii**). Finally, two-dimensional Gaussian mixture models (GMMs) fitted to median and standard deviation features confirmed this separation, yielding a Jensen-Shannon (JS) divergence of 0.48 (Figure 7C**.iv**). The combination of these metrics strongly indicated that the *S. cerevisiae* tRNA isodecoders Arg-ACG-1-1 and Arg-ACG-2-1 have unique ionic current profiles. Applying the same comparison strategy on an example isodecoder pair in *E. coli* showed distinct populations of Thr-GGT isodecoder variants. (**Supplementary Figure 8**).

When we compared the Gaussian mixture models across all isoacceptors with isodecoders for both *S. cerevisiae* and *E. coli*, the JS divergence ranged from 0.28 to 0.98 (**Supplementary Figures 9 and 10**, **Supplementary Table 9**). For *S. cerevisiae,* the isodecoder pair Ser-AGA-1-1 and Ser-AGA-2-1 had the lowest JS divergence at 0.28. This can possibly be attributed to the nature of the sequence level divergence between the two isodecoders: Ser-AGA-2-1 has a dinucleotide GG spanning consecutive Sprinzl coordinates V4, V11, where Ser-AGA-1-1 lacks the V4 position entirely, meaning it only has a G mononucleotide at V11. This means that the entire sequence difference between the two isodecoders is GG for Ser-AGA-2-1 and G for Ser-AGA-1-1^34,36^. For *E. coli*, the fMet-CAT-1-1 and fMet-CAT-2-1 had the lowest JS divergence at 0.37. fMet-CAT-1-1 and fMet-CAT-2-1 have a Guanosine and Adenosine at position 46, respectively. According to MODOMICS^48^ fMet-CAT-1-1 can have an m^7^G at position 46, potentially further complicating discrimination of the two isodecoders.

While these two-dimensional summaries provide intuitive insight into population-level differences, they capture only a fraction of the underlying ionic current signal complexity. The classification model leverages the full temporal structure of ionic current data, incorporating dependencies across time points that median and standard deviation amplitude alone cannot represent. These results suggested that tRNAZAP could separate out two distinct ionic current populations associated with isodecoders. Additionally, tRNAZAP achieved higher alignment identity. This suggests that tRNAZAP is successfully resolving isodecoders in the absence of ground truth biological labels.

### tRNAZAP captures abundance changes in biological *S. cerevisiae* tRNA

Nearly all eukaryotic organisms harbor tRNA genes in two different genomes: the nuclear genome and the mitochondrial genome. *S. cerevisiae* grown in glucose-based media utilize fermentation as a primary energy source leading to decreased mitochondrial abundance. This results in fewer mitochondria and by extension fewer mitochondrial tRNA^49^. The abundance of mitochondrial tRNA can be increased in a sample by using different growth media and an enrichment of the mitochondrial fraction^44,50–52^. We wanted to test if tRNAZAP could detect the enrichment of an entire subpopulation of tRNA simultaneously. To that end, we sequenced tRNA from a mitochondria-enriched tRNA sample (MT+, see methods). We compared the MT+ condition with the glucose growth condition (MT-) and documented a 4.1-to-8.7 Log_2_ FC for the 24 *S. cerevisiae* mitochondrial tRNA isodecoders, demonstrating tRNAZAP’s ability to capture subpopulation enrichment (Figure 8). We noted that the median alignment identity for mitochondrial tRNA from the enriched condition was equal to or greater than mitochondrial tRNA from the yeast grown in glucose across all isodecoders (**Supplementary Table 10**).

**Figure 8.**
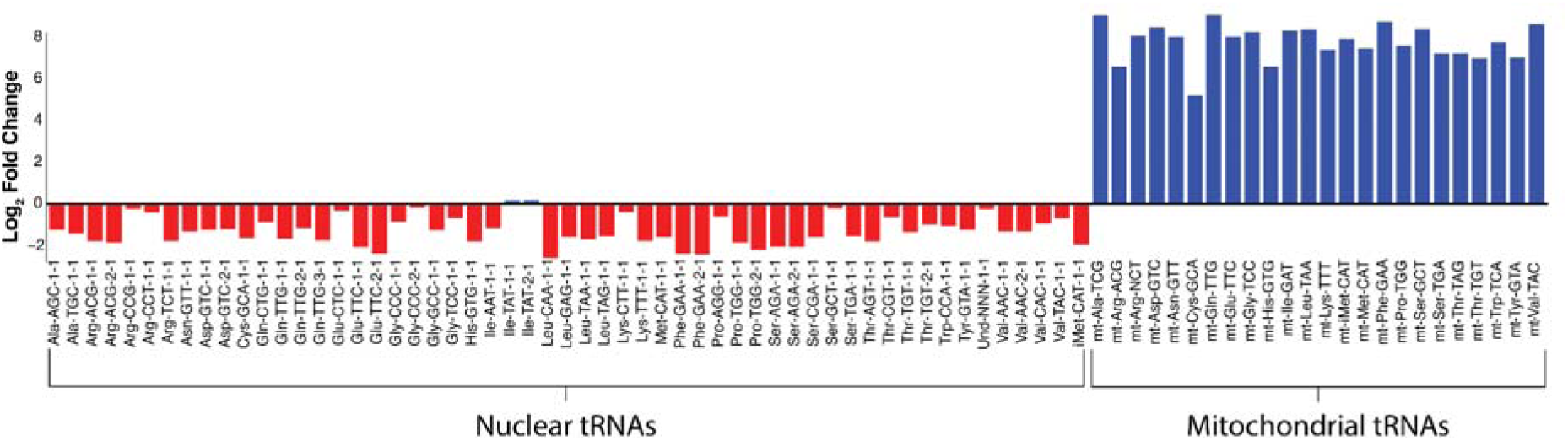
Applying tRNAZAP to enriched biological *S. cerevisiae* tRNA. Log_2_ Fold Change between the TPMs of the mitochondrial-enriched condition and the non-enriched condition. The mitochondrial enrichment condition is the numerator and wild wild-type condition is the denominator when calculating log_2_ fold change. The x-axis shows each of the tRNA isodecoders in the reference set for tRNAZAP while the y-axis shows the log_2_ fold change. Nuclear tRNAs and mitochondrial tRNAs are delineated with brackets and a label.

### Nucleotide modification analysis using ionic current data

In previous work, we and others investigated the modification circuit of the T-loop in *S. cerevisiae* tRNA^6,12,53^. We combined mass-spectrometry, genetic controls, and direct RNA sequencing to analyze the inter-dependence of T-loop modifications. Because tRNAZAP model was trained using wild-type *S. cerevisiae* Nanopore tRNA data, we wanted to test its performance on Nanopore tRNA data from a genetically altered *S. cerevisiae* strain. We sequenced total tRNA from *pus*4Δ *S. cerevisiae* strain. This strain lacks the pseudouridine synthase gene Pus4, that modifies U55 on nearly all *S. cerevisiae* tRNAs^6^. tRNAZAP aligned more reads than BWA-MEM for the *pus*4Δ strain. Additionally, tRNAZAP had a higher alignment identity than BWA-MEM for reads aligned by both aligners and reads where the aligners disagreed (**Supplementary Figure 11**).

This increase in number of aligned reads coupled with higher alignment identity lends itself to investigations of ionic current perturbations caused by putative modifications. While miscalls have provided suggestive evidence in tRNA modification analysis using Nanopore DRS, it is important to adopt principled approaches using ionic current as the community continues to research tRNA modifications. To qualitatively recapitulate the miscall-based findings of our previous work, we investigated the ionic current profiles for the T-loop in *S. cerevisiae* tRNA. We examined Cys-CGA-1-1 because it bears a 5-methyluridine (m^5^U) at position 54, pseudouridine (Ψ) at position 55, and a 1-methyladenosine (m^1^A) at position 58^48^. This cluster of modifications produced a distinct miscall pattern. In our prior work, we had documented that the miscall signal for both Ψ55 and m^1^A58 reduced significantly in tRNA data for the *pus*4Δ strain^6^ (Figure 9A). Nucleoside LC-MS had confirmed that all three modifications in the T-loop were reduced when Pus4 was knocked out. This work was performed using the older RNA002 platform that had lower throughput, lower basecalling accuracy, and at the time of analysis no access to the moves table.

**Figure 9.**
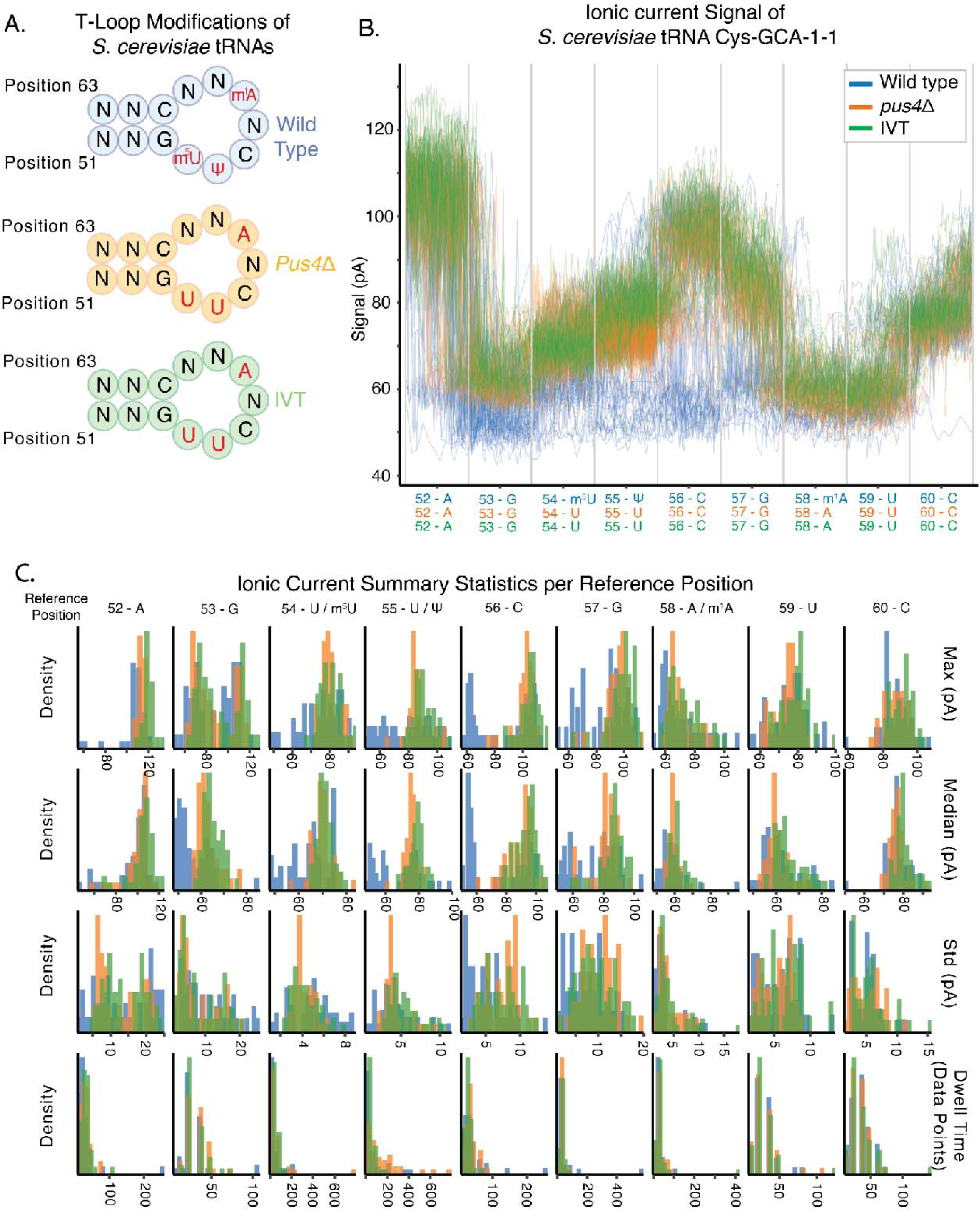
Ionic current signal comparison of tRNA Cys-GCA across conditions with *pus*4Δ. **A.** Illustration of the modified positions in the T-Loop of *S. cerevisiae* across three conditions: wild-type (blue); *pus4*Δ (orange), a knockout of the *pus4* enzyme which is responsible for the addition of the pseudouridine at position 55; and in vitro transcribed (green) tRNAs which completely lack modifications. **B.** Representative ionic current traces for tRNA Cys-GCA across three experimental conditions: Wild type (blue), *pus4*Δ (orange), and IVT (green), each with ∼50 reads. Signals are displayed in picoamperes (pA) and tRNA sequence positions (51-63). Traces are visualized based on the Dorado moves table (a signal-to-read alignment that maps blocks of raw signal timepoints to basecalled nucleotides. The x-axis indicates genomic position and corresponding nucleotide identity). **C.** Summary statistics per position showing distributions of signal characteristics across the three conditions. Each column represents a nucleotide position with three statistical measures displayed as density plots: median signal intensity (Median, top row), standard deviation of signal intensity (Std, second row), and signal dwell time in timepoints (Duration, bottom row). Density curves are color-coded as in panel A, with shaded areas representing the distribution of per-read values for each condition at each position.

Using the RNA004 DRS platform, we examined ionic current traces for Cys-CGA-1-1 tRNA T-loop region from the wild type condition, the IVT condition, and the Pus4 knockout condition (Figure 9B). We documented two distinct populations in the ionic current in the vicinity of the modification-bearing positions in the T-loop. We plotted the position-wise histograms of maximum picoamperes (pA), median pA, standard deviation, and duration (Figure 9C). The IVT and *pus*4Δ ionic current traces and histograms showed concordance throughout the region. This follows the documented modification circuit suggesting *pus*4Δ reads that lack Ψ55 would additionally have reduced levels of m^1^A58 and m^5^U54. The wild type condition also has some reads that follow the same ionic current pattern observed in the IVT and *pus*4Δ populations suggesting a similar modification profile. However, the wild type condition has a second population of reads that produce both a unique ionic current pattern observed in Figure 9B positions 53-57 and distinct lower peaks in median picoamperes in Figure 9C positions 53-57. A possible explanation for this, is that the wild type specific patterns of ionic current are for strands that have the complete or partial set of tRNA Cys-GCA-1-1 T-loop modifications: m^5^U54, Ψ55, and m^1^A58. It is important to note that the nanopore sensor reads a series of consecutive nucleotides in an RNA strand at a time (typically modeled as a 5mer), moving in a single nucleotide stepping pattern. Thus, the influence of a nucleotide modification can extend beyond the nucleotide position^18,25^. In the wild type condition, we observed approximately 5 positions of impacted signal. This could be due to the combined influence of ionic current from the trio of modifications m^5^U54, Ψ55, and m^1^A58 spread across the observed positions. The separation of populations within the wild-type condition suggests that with proper training, controls, and orthogonal validation, ionic current analysis could document modification stoichiometry in tRNA.

## DISCUSSION

We demonstrated that tRNAZAP outperformed sequence-based techniques for the analysis of tRNA isodecoders from *E. coli* and *S. cerevisiae* by aligning more reads with higher alignment identity. Ionic current data-based models captured the expected populations in enriched tRNA samples (tRNA^Phe^ in *E. coli* and mitochondrial tRNA in *S. cerevisiae*) demonstrating broad applicability across growth and purification conditions. Additionally, tRNAZAP classification documented distinct ionic current distributions between closely related tRNA isodecoders. The increased read count and alignment identity of tRNAZAP permitted RNA modification analysis based on ionic current when comparing wild-type and *pus*4Δ knockout conditions for *S. cerevisiae*.

A present requirement of tRNAZAP is the use of specific synthetic adapters. tRNAZAP considers the entire ionic current readout including these adapters, meaning nanopore tRNA data generated using other splint adapters could behave unpredictably. We believe that the increased throughput coupled with the higher quality of data for each experiment justify these tRNAZAP adapters. We recognize that tRNAZAP isodecoder assignments have not been validated with molecular biology techniques. However, this would require purification of individual tRNA isodecoders which is challenging for all contemporary separation techniques such as Chaplet Column Chromatography. Better purification strategies are required.

We anticipate that nanopore ionic current will be a key component of the principled approaches to study tRNA isodecoders and modifications. The combination of nanopore direct tRNA sequencing coupled with orthogonal validation of LC-MS/MS provides a strong foundation and standard for tRNA research. The success of tRNAZAP for *E. coli* and *S. cerevisiae* is predictive of its potential for more complex tRNA populations. This will be foundational for studying the comprehensive landscape of human tRNA isodecoders and their modification interplay.

## METHODS

All of the adapter oligonucleotides, IVT oligonucleotides and DNA ultramer pools used in these studies were manufactured by IDT. The adapter oligonucleotides were HPLC purified,

The biological *E. coli* tRNA sample sequenced was from MRE600 total tRNA purchased from Roche Pharmaceuticals. The *S. cerevisiae* samples were from strain BY4741 collected during log phase growth.

### Enrichment of tRNA-Phe from *E. coli* total RNA

Wild-type E. coli K-12 BW25113 were transformed with a pUC57 plasmid containing a lac-inducable tRNA-Phe-GAA-1-1 gene for overexpression, then struck onto LB-amp plates and grown overnight at 37°C. Individual colonies were selected and grown to saturation in LB with ampicillin. An expression culture was inoculated at a 5:1000 ratio in enriched TB with ampicillin, 0.05% glucose, and 0.2% lactose for autoinduction, and then grown for 16 hours before harvesting in a centrifuge. Pseudopurified tRNA phe was generated as previously described^54^. Briefly, cells were lysed and total RNA was isolated using a phenol chloroform extraction. Tris-acetate pH 8 was used to deacylate the tRNA in the sample and then total tRNA was isolated using a Cytiva Resource Q anion exchange column on an FPLC. This total tRNA was ethanol precipitated and a Waters XBridge BEH C18 OBD prep HPLC column was used for pseudopurification. Fractions containing high tRNA phe concentrations were identified using a test charging with radiolabeled phenylalanine and pooled.

### *S. cerevisiae* wild type growth conditions

Wild-type BY4741 *S. cerevisiae* were struck out from glycerol stocks onto YPD plates and grown for three days at 30°C. Single colonies were inoculated in 3mL liquid YPD and grown overnight at 30°C, after which 400uL of overnight culture were diluted into 5.2mL liquid YPD and grown at 30°C to an OD of 0.5-1.0. Cell pellets were washed with 1X PBS and flash frozen in liquid nitrogen. Total RNA was extracted using the hot phenol method as previously described^6^.

### *S. cerevisiae* mitochondrial growth conditions and mitochondrial enrichment

Wild-type BY4741 *S. cerevisiae* were grown in 1 L YPE (Yeast extract, Peptone, 2% Ethanol) and flash frozen in liquid nitrogen at an OD600 of 1.0-1.4. Differential centrifugation was used to isolate the crude mitochondrial fraction from *S. cerevisiae* cells as previously described^44^. RNA was isolated from the crude mitochondrial fraction using the hot phenol-chloroform method.

### Biological tRNA adapter augmentation for deep learning

Previous work by our group and others share the common theme of adapting the tRNA by ligating 5’ and 3’ adapter strands to the conserved NCCA tRNA 3’ overhangs^12,16^. To facilitate model learning we amended our original adapter lengths to have approximately equal lengths of sequence bearing constructive ionic current information. Our final constructs had a 5’ adapter length of 36 and 3’ length of 60, of which 10 are poly(A) and another 18 are the DNA nucleotides required for the customized RTA ligation. Here we consider the 32 nucleotides before the poly(A) to approximately equal in length to the 5’ adapter. The 5’ adapter was designed to complement each of the 5 known 3’ tRNA overhangs (ACCA, UCCA, GCCA, CCCA, CCA). The 3’ adapter included 18 DNA nucleotides for the addition of the ONT SQK-RNA004 sequencing adapter and a poly(A) homopolymer for demarcation of the start of the 3’ RNA sequence. The sequences for these oligos can be found in **Supplementary Table 2**.

### Candidate tRNA Reference Curation

For *S. cerevisiae*( https://gtrnadb.org/genomes/eukaryota/Scere3/sacCer3-mature-tRNAs.fa) and *Escherichia coli* (https://gtrnadb.org/genomes/bacteria/Esch_coli_K_12_MG1655/eschColi_K_12_MG1655-mature-tRNAs.fa) we downloaded the set of genomically predicted tRNA sequences from gtRNA-db. The set of unique sequences were curated for each species and the sequences were attributed to the first predicted tRNA of that sequence based on tRNA naming conventions ^34^. This set of tRNAs represents a comprehensive set of possible tRNA sequences based on genomic predictions. In our previous work a more restrictive iso-acceptor level reference was curated to account for the difficulties of resolving minimal sequence differences using a seed and chain algorithm strategy such as BWA-MEM. These sequences were used as references for alignment using tRNAZAP (**Supplementary Table 1).**

### Design of tRNA DNA Templates for *in vitro* transcription

We appended the 3’ adapter and the appropriate 5’ adapter to the corresponding side of each target tRNA sequence. The IVT template is the reverse complement of this unified sequence with an additional T7 binding site sequence (TATAGTGAGTCGTATTAAATGATG) included on the 3’ most end of each template. From the complete set of template sequences the interspecies and intraspecies pairwise sequence similarity was calculated using levenshtein distance (edit distance) as the score. We pooled the *E. coli* and *S. cerevisiae* tRNA sequences and split them into 8 groups that allowed a minimum similarity score of 16, where lower scores represented more similar sequences. Each group of sequences was ordered as an Oligo pool from Integrated DNA Technologies. We prepared a custom reference for the merged *E. coli* and *S. cerevisiae* IVT products (**Supplementary Table 11**)

### *In Vitro* Transcription

The 130 IVT templates were pooled in 10ul (162.5pmol total) and annealed with a two fold molar amount (325pmol) of the T7 oligonucleotide in 1X TNE by heating to 75C for 1 min followed by slow cooling. One microgram of the annealed oligo/template mix, was used to produce our tRNA representative RNA using the NEB HiScribe T7 Quick High Yield RNA Synthesis Kit (NEB Catalog #E2050) following manufacturer’s recommendations for standard RNA synthesis of short RNA (<300nt). The reactions went for 16 hours, then were DNAse I treated for 30 minutes and cleaned up using 1.5X RNACleanXP beads (Beckman-Coulter) according to manufacturer’s protocol and eluted in NF H20.

### *In Vitro* Transcription Library Preparation and Sequencing

For each experiment an augmented SQK-RNA004 (Oxford Nanopore technology)protocol was followed that used a custom splint adapter in the place of the RTA adapter. The first ligation was composed of 2 ug of the purified in vitro transcription reaction, 6µl of 5x quick ligation buffer (NEB Catalog #B6058S), 34 pmol of the custom adapter and 3 µl of T4 DNA ligase (2,000,000 units/mL, NEB Catalog #M0239S) in a total volume of 30ul. The protocol was followed from this point forward. Sequencing took place on two separate devices; a PromethION48, and PromethION2.

### Biological Library preparation and Sequencing

RNA Ligase II (NEB) was used to ligate tRNA to the five splint adapters RNA. The reactions typically had 8-10 pmol of deacylated tRNA and a 1.1 to 4 fold excess of adapters. Further detail of the ligation conditions for different runs are shown in **Supplementary Table 12**. The ligation reactions went for 45-60min at RT and were cleaned up with 1.3X Ampure RNACleanXP Spri beads and eluted in 23ul of NFH20. The eluate was then ligated to 4.5ul RLA motor associated adapter using T4 DNA ligase (2,000,000 U/mL) for 30 min and cleaned up using 0.8Xbeads, washed with WSB and eluted in 33ul REB. PromethION flow cell priming and sequencing followed the RNASQK004(version X) and were run with basecalling off.

### Basecalling and BWA-MEM Alignment

All experiments were basecalled with Dorado 1.0.0 model rna004_130bps_sup@v5.2.0. For standard alignment each experiment was aligned to a custom isoacceptor level reference that included the sequencing adapters and using BWA-MEM. Tags bearing ionic current and training relevant information were transferred from the unmapped reads to aligned reads using pysam ^55^.

### Data Labeling for IVT training

Data were aligned to our complete custom reference sequence set (**Supplementary Table 13**) representing the expected isoacceptor level IVT product using BWA-MEM.

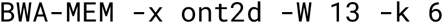

Reads with a mapping quality of 0 were removed and reads were labeled according to their primary tRNA alignment. An additional round of filtering removed:

1. Reads with less than 15 positions of aligned tRNA sequence,
2. Reads with lower than 75% identity across the entire reference span including splints

In addition to labeling the isodecoder of each read, sections of signal were labeled as the DNA adapter, 3’ Adapter, tRNA of interest, and 5’ Adapter by assigning bases based on the corresponding Dorado moves table signal to base assignments.

The produced data and their corresponding labels were fed into the model training pipeline.

### tRNAZAP Inference Pipeline

Our pipeline processes nanopore ionic current signals through two integrated components: (1) the classification and segmentation model and (2) the Aligner.

### Classification and Segmentation model

Our deep learning model processes raw nanopore ionic current signals to perform two key functions for tRNA analysis:

- **Isodecoder-level Classification:** Determines the specific tRNA isodecoder present in the signal.
- **Ionic Current Segmentation:** Isolates the tRNA-containing region by identifying and removing ONT adapters and 3’/5’ splints.

### Model Architecture

We implement these functions using a multitask Transformer neural network that divides the continuous ionic current into discrete, non-overlapping time windows (chunks). This chunking approach enables the model to learn both local patterns within individual windows and global dependencies across the entire signal. A *[CLS]* token is prepended to the input sequence, whose embedding is used for isodecoder classification via a dedicated classification head. The embeddings corresponding to each signal chunk are independently classified by another classification head into one of four possible regions.

### Model Training and fine-tuning

The multitask transformer was trained jointly on the classification and segmentation objectives using the labels from section *Data Labeling*. The total loss was computed as a weighted combination of two categorical cross-entropy losses (for both isodecoder-level classification and ionic current segmentation) with weights of 0.3 and 0.7, respectively. Models were optimized using the Adam optimizer with learning rate scheduling. Training was performed for 50 epochs with a base learning rate of 0.001. During finetuning, Model parameters were updated using a learning rate of 0.0001 over 30 epochs.

### Evaluation Metrics

Performance was evaluated with specific metrics for each task by comparing predicted labels versus the labels from section *Data Labeling*.

For sequence classification, we report micro and macro accuracy. Micro accuracy reflects overall performance across all reads, while macro accuracy averages performance across isodecoders to account for class imbalance.

For segmentation, we report the mean Intersection-over-Union (IoU) of the predicted versus BWA-MEM tRNA region across the dataset.

For each sequence *i*, the IoU is computed between the predicted and BWA-assigned tRNA regions (defined by start and end positions of the tRNA region). Let *P_i_* denote the predicted region and *L_i_* the BWA-MEM-assigned region. The IoU is given by:

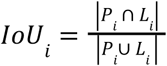

where |*P_i_* ∩ *L_i_*| is the length of the overlapping region (Intersection of regions) and |*P_i_* ∪ *L_i_*| is the length of the union of the two regions. A visual example is provided in **Supplementary Figure 11**.

The overall segmentation score is then reported as the average across all L sequences:

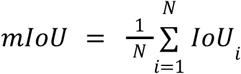

### Model Polishing with Biological Data, Data Labeling

For both *E. coli* and *S. cerevisiae*, we used biological sequencing runs to fine-tune the model to be cognizant of tRNA-specific modification profiles in ionic current space. To disambiguate isodecoders from biological data we aligned to the isoacceptor level reference and created a lookup table for all of the isodecoder level variants. For a read to count towards an isodecoder, it had to have a minimum alignment identity of 75% and have a positive match at the correct position for every isodecoder-specific variant. In the case of *S. cerevisiae* Asp-GTC and Ser-AGA which have isodecoder with different lengths, we assigned isodecoder identity by a Needleman-Wunsch edit distance calculation. The isodecoder requiring fewer edits was selected for fine-tuning. Reads that could not fulfill the overall identity, isodecoder-specific identity, or edit distance requirements were not used in fine-tuning.

### Model Polishing with Biological Data, Training

For fine-tuning the IVT models on biological data, a collection of biological tRNA were first collected. Biological data exhibit isodecoder imbalance due to rarity of some of the tRNA types in nature. To address this imbalance, we randomly sampled 200,000 examples per isodecoder for fine-tuning. isodecoders with fewer than 20,000 samples retained all available examples. This procedure reduces the overrepresentation of highly abundant isodecoders while keeping the population of biologically rare tRNA types limited. The models were allowed to train on the data for 30 epochs. Performance was evaluated with the same metrics for IVT model performance.

### tRNAZAP Aligner

tRNAZAP aligner uses the predicted tRNA isodecoder, ionic current boundaries for the tRNA, and the moves table produced by Dorado to align to an isodecoder-specific tRNA reference with an augmented Needleman-Wunsch algorithm. The algorithm finds the path traversing the entire tRNA reference for the predicted isodecoder that maximizes the ratio of query bases explained over edit distance. Optionally, the aligner can consider the highest probability isodecoder or the second highest probability isodecoder, ultimately selecting the option with the lower edit distance. For possibly fragmented reads a check is performed: if 15% of the reference length is accounted for by deletions an affine-gap Smith-Waterman alignment is performed to find a sub alignment between the query and the reference. Finally, an optional rescue alignment can be performed if the tRNA region alignment fails. The last shot alignment performs a Smith-Waterman alignment of the entire query sequence against the top 3 tRNA references. A minimum of 25 matches in the tRNA sequence is required as well as a minimum identity of 75%. These alignments are output in a SAM compliant format and preserve all of the original reads SAM Tags.

### Statistical analysis of isodecoder signal differences

For each isodecoder pair, we compared ionic current signal features extracted from a fixed window centered on the site of sequence difference. For each read, two summary features were computed: the median ionic current and the standard deviation of the ionic current within the window. Thus, isodecoder pairs L and L are represented with two-dimensional feature vectors 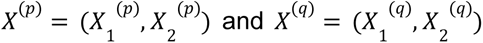, where *X*_1_ is the vector of per-read medians and *X*_2_ is the vector of per-read standard deviations of the ionic currents measured within the window.

For each feature *i* ∊ {1,2}, we tested the marginal null hypothesis

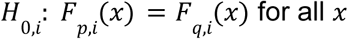

where *F_p,i_* and *F_q,i_* denote the cumulative distribution functions of feature L for the two isodecoders. The corresponding alternative hypothesis was

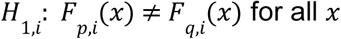

Marginal hypotheses were evaluated using two-sample Kolmogorov–Smirnov tests. To control the family-wise error rate across the two tested features, *p*-values were adjusted using the Bonferroni correction. The family-wise significance level was set to α = 0.05, corresponding to a per-feature significance level of α/2 = 0.025.

The family-wise null hypothesis, that all marginal feature distributions are identical between isodecoders, was rejected if any Bonferroni-adjusted p-value was below α.

To summarize differences in the joint distribution of the two features, we additionally computed the Jensen–Shannon (JS) divergence between two-dimensional Gaussian mixture models fitted to the feature distributions of each isodecoder. JS divergence was used as a descriptive measure of joint distributional differences and was not used for formal hypothesis testing.

## Data Availability

Sequence-level data will be publicly available through the European Nucleotide Archive. Signal files are available on request from the authors. tRNAZAP is available at GitHub repository (https://github.com/genometechlab/tRNA_zap).

## Acknowledgements

The authors would like to acknowledge Joseph Roffi and the ISEC-Ops team for maintaining the facilities and promoting a healthy working environment.

## Author Contributions

S.A., T.T., J.L.R., M.L.B. M.Z., R.A.S. contributed to experimental work. S.A. and P.D.K. contributed to data analysis, writing, and editing. N.G.B. contributed to data analysis, writing, and editing. D.M.G. and K.S.K. contributed to experimental work, writing and editing. M.J contributed to supervision, conceptualization, experimental work, writing, and editing.

