## Supplementary Figures for "tRNA isodecoder analysis using Nanopore ionic current signals and deep learning"

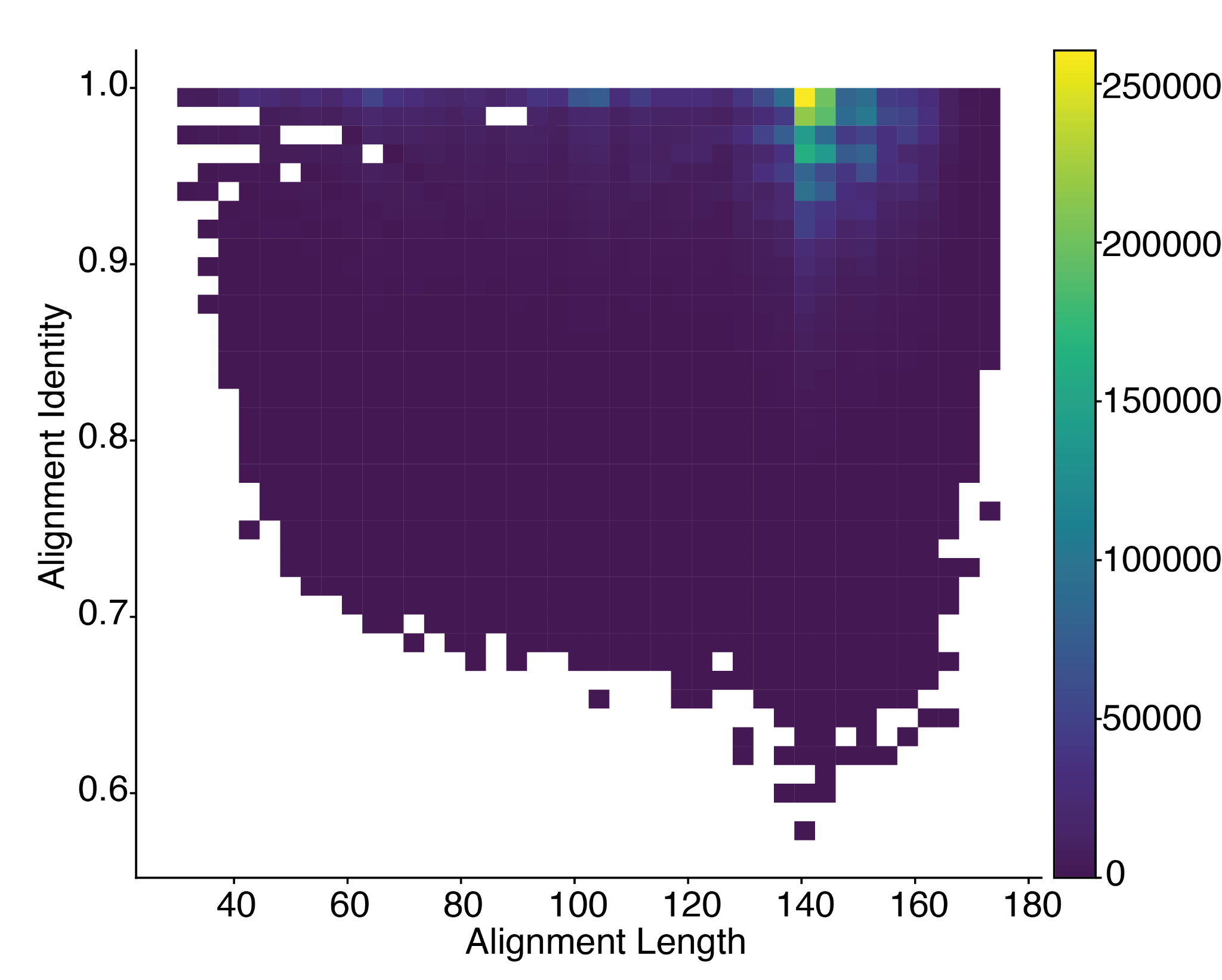

**Supplementary Figure 1. 2D Distribution of alignment length and alignment identity for IVT tRNA.** The x-axis represents the aligned length of IVT tRNA reads using BWA-MEM and aligning to the reference set in **Supplementary Table 9**. The y-axis reports the alignment identity. The color of each cell describes the number of reads that fell in that combination of alignment length and identity.

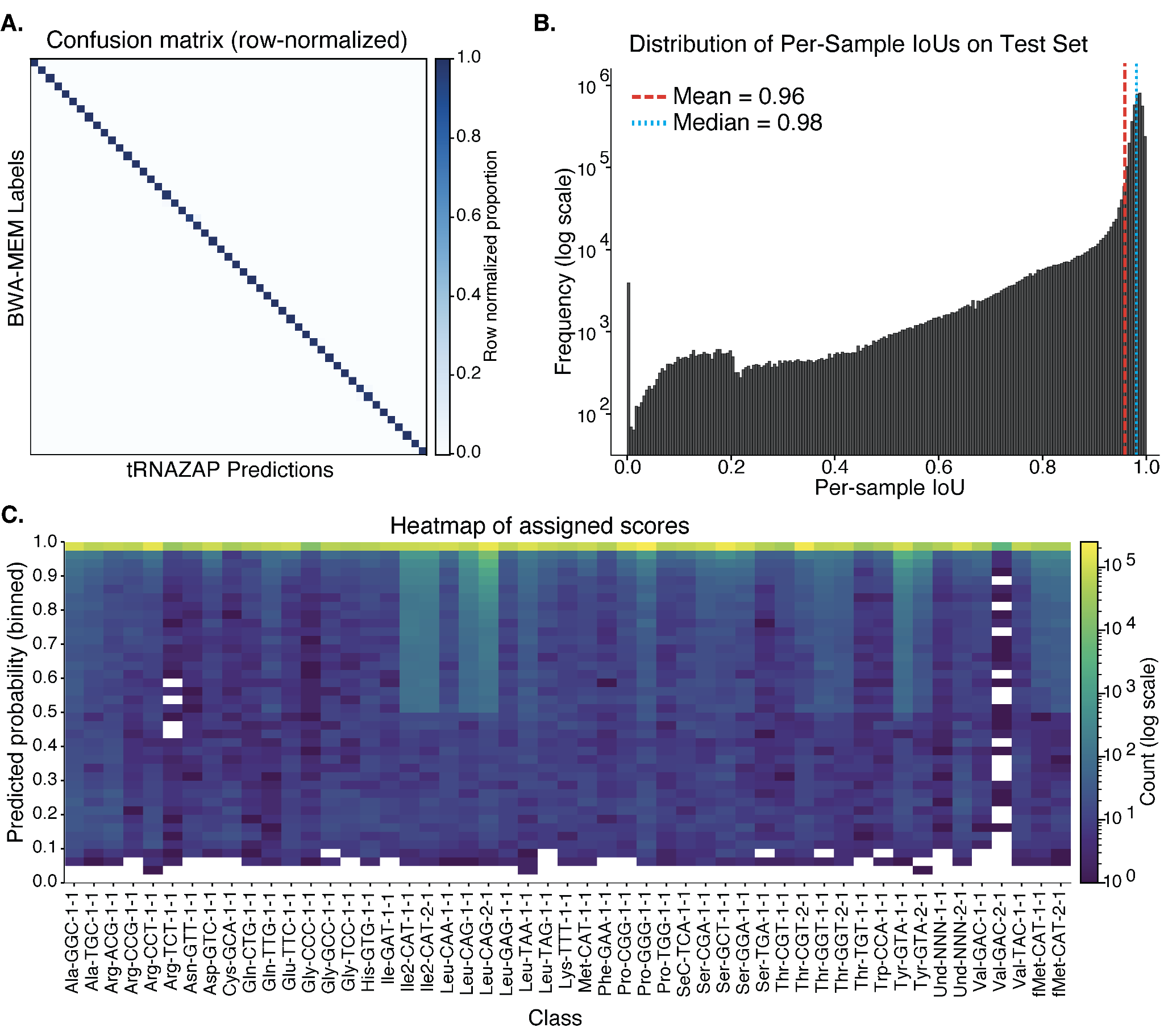

**Supplementary Figure 2.** tRNAZAP’s inference model performance on the test subset of *E. coli* IVT reads when compared to BWA-MEM based sequence level classification as well as the Dorado move table’s signal to sequence mapping output. **A.** Confusion matrix illustrating the correspondence between BWA-MEM-derived reference labels and the model’s final classifications. BWA-MEM reference labels were enforced to have a minimum of 75% identity. **B.** Distribution of Intersection-over-Union (IoU) scores between the model-predicted segment ranges and the BWA-MEM assigned labels. The Intersection-over-Union metric allows for the mathematical analysis of overlap between BWA-MEM and move table based segmentation of the tRNA region against the tRNAZAP inference based segmentation of the inference region.

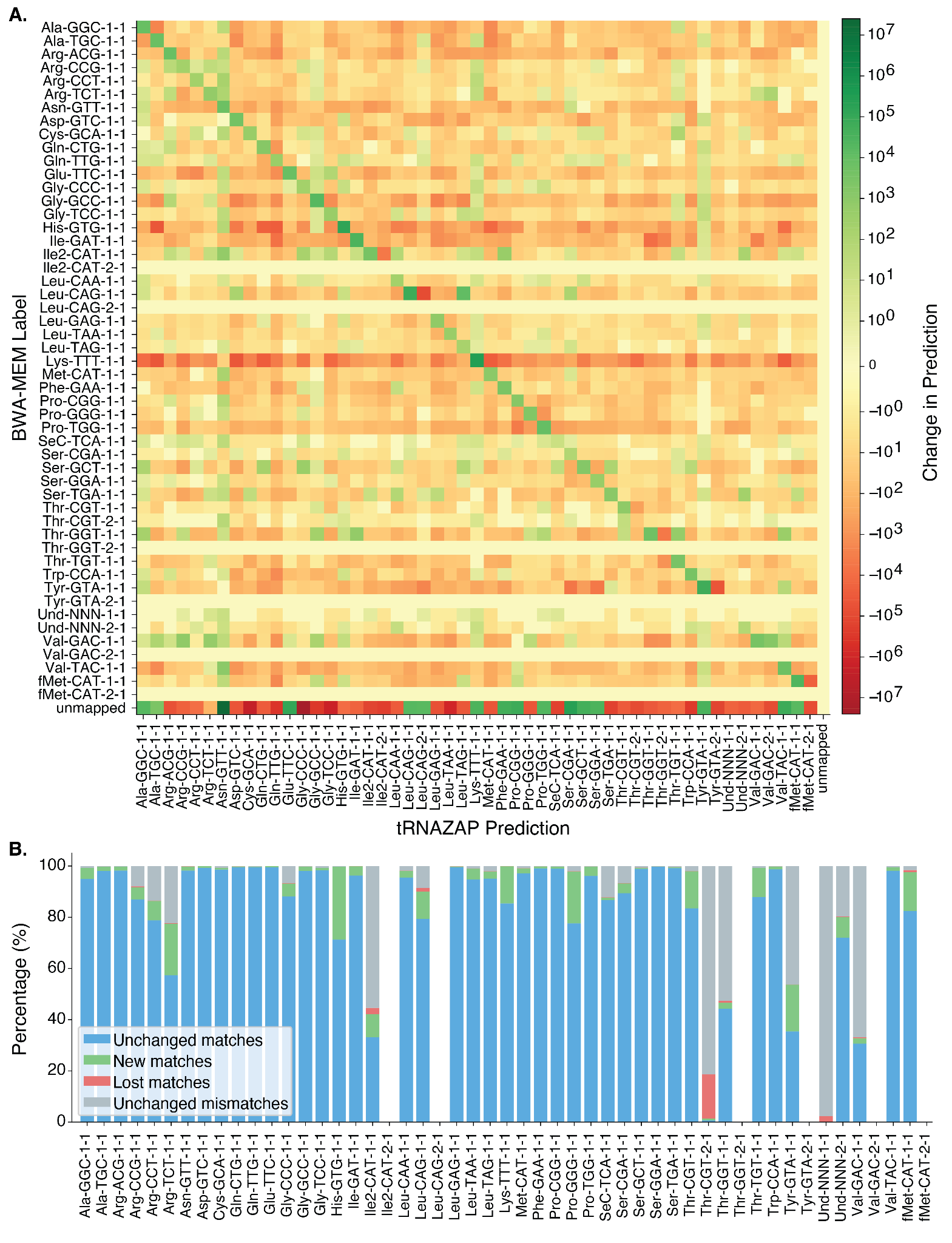

**Supplementary Figure 3.** Effect of model fine-tuning on performance for biological data. **A.** Heatmap illustrating changes in tRNA isodecoder assignments following biological fine-tuning. Rows correspond to BWA-MEM assigned labels, while columns represent tRNAZAP model predictions. The color scale indicates the log-count of samples per cell. Increased cell values along the diagonal demonstrates improved agreement between tRNAZAP predictions and BWA-MEM labels after fine-tuning. The rows composed entirely of 0s come from tRNA isodecoders that are withheld from the reference to accommodate the BWA-MEM alignment scoring system. The only exception is the Isoacceptor Thr-CGT-2-1 which has sufficient sequence level differences from Thr-CGT-1-1 to minimize alignment impact. **B.** Class-wise redistribution of tRNAZAP prediction outcomes following fine-tuning. For each isodecoder defined by the BWA-MEM reference alignment, the stacked bars show the percentage of reads grouped according to how tRNAZAP’s predicted isodecoder changes with fine-tuning. Blue indicates reads for which the predicted isodecoder matches the BWA-MEM label both before and after fine-tuning (unchanged matches). Green denotes reads whose predicted isodecoder differs before fine-tuning but matches the BWA-MEM label after fine-tuning (new matches). Red represents reads whose predicted isodecoder matches the BWA-MEM label before fine-tuning but differs after fine-tuning (lost matches). Gray corresponds to reads whose predicted isodecoder differs from the BWA-MEM label both before and after fine-tuning (unchanged mismatches). Percentages are computed independently within each BWA-MEM-defined isodecoder and normalized to 100% for each bar.

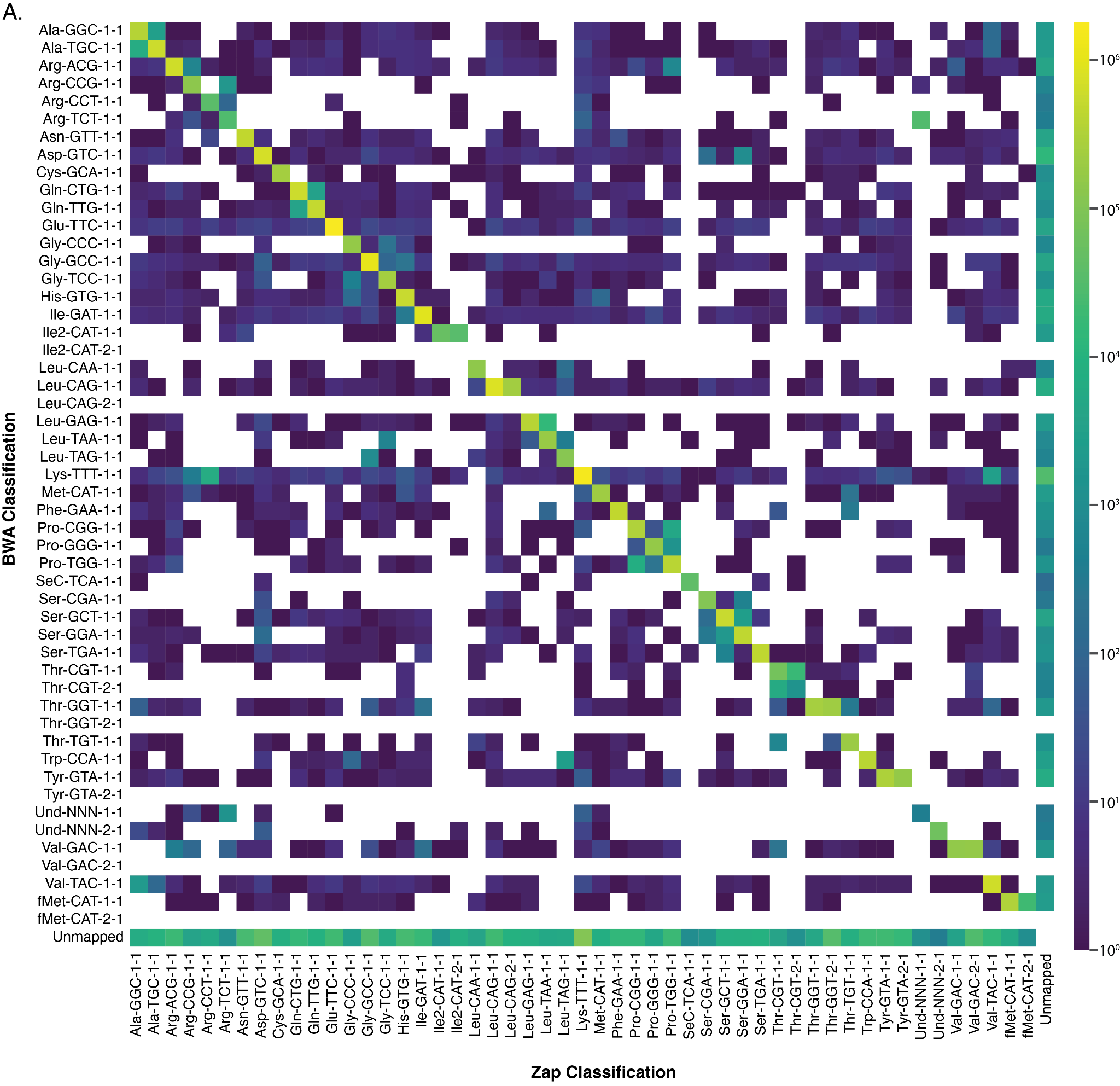

**Supplementary Figure 4.** Heatmap showing the aligned read counts for tRNAZAP and BWA-MEM across all *E. coli* isodecoders. The x-axis shows the reference that tRNZAP aligned the read to, while the y-axis shows the reference that BWA-MEM aligned the read to. The color of each cell represents the total number of reads that had that specific combination of tRNAzap reference and BWA-MEM reference.

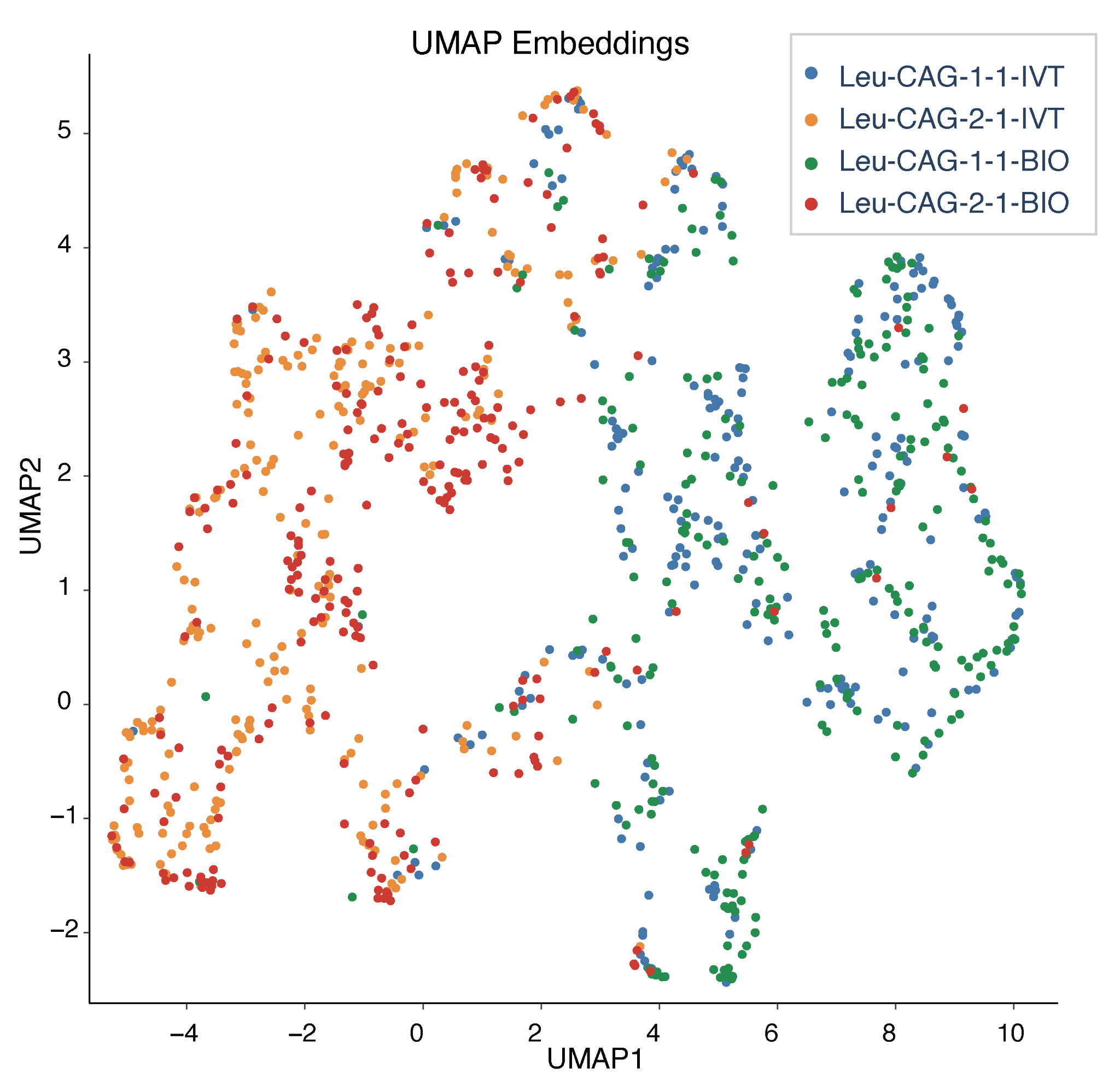

**Supplementary Figure 5. UMAP of E. coli tRNA isodecoder variants in biological and IVT conditions.** UMAP projection of multi-position ionic current features, revealing population structure between IVT and biological isodecoders. We took the median pA and standard deviation of each position from 45 to 47, for Leu-CAG isodecoder variants from biological and IVT E. coli. The UMAP coloring matched the isodecor names.

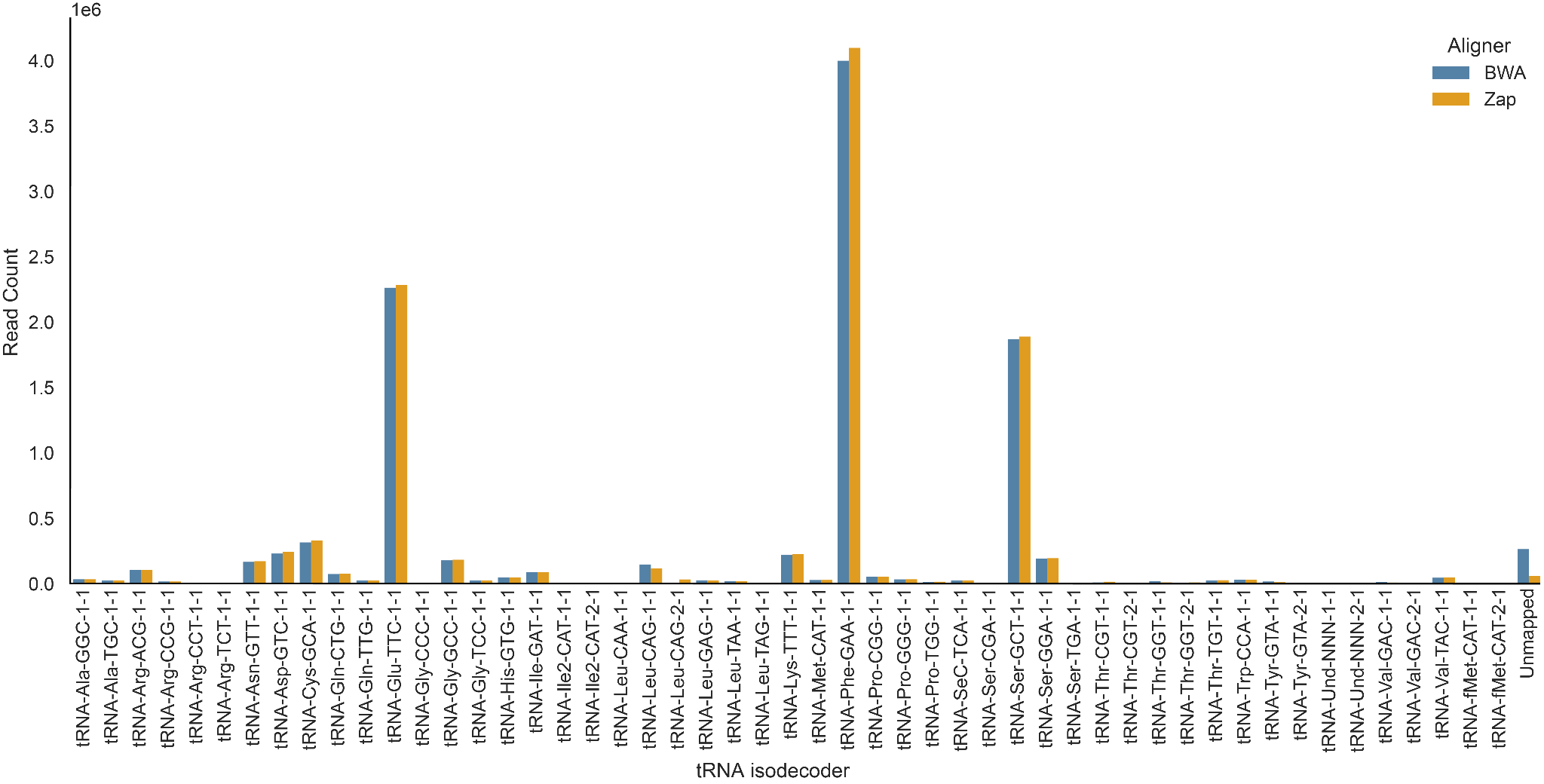

**Supplementary Figure 6. BWA-MEM Co-enrichment populations for *E. coli* tRNA Phe targeted enrichment.** The x-axis shows the different *E. coli* tRNA isodecoders present in the reference set that tRNAZAP is aligned to. The same reference set was available to BWA-MEM barring the isodecoders which are delineated with a 2-1 in the x-axis labels. The y-axis shows the count of reads aligned to each class.

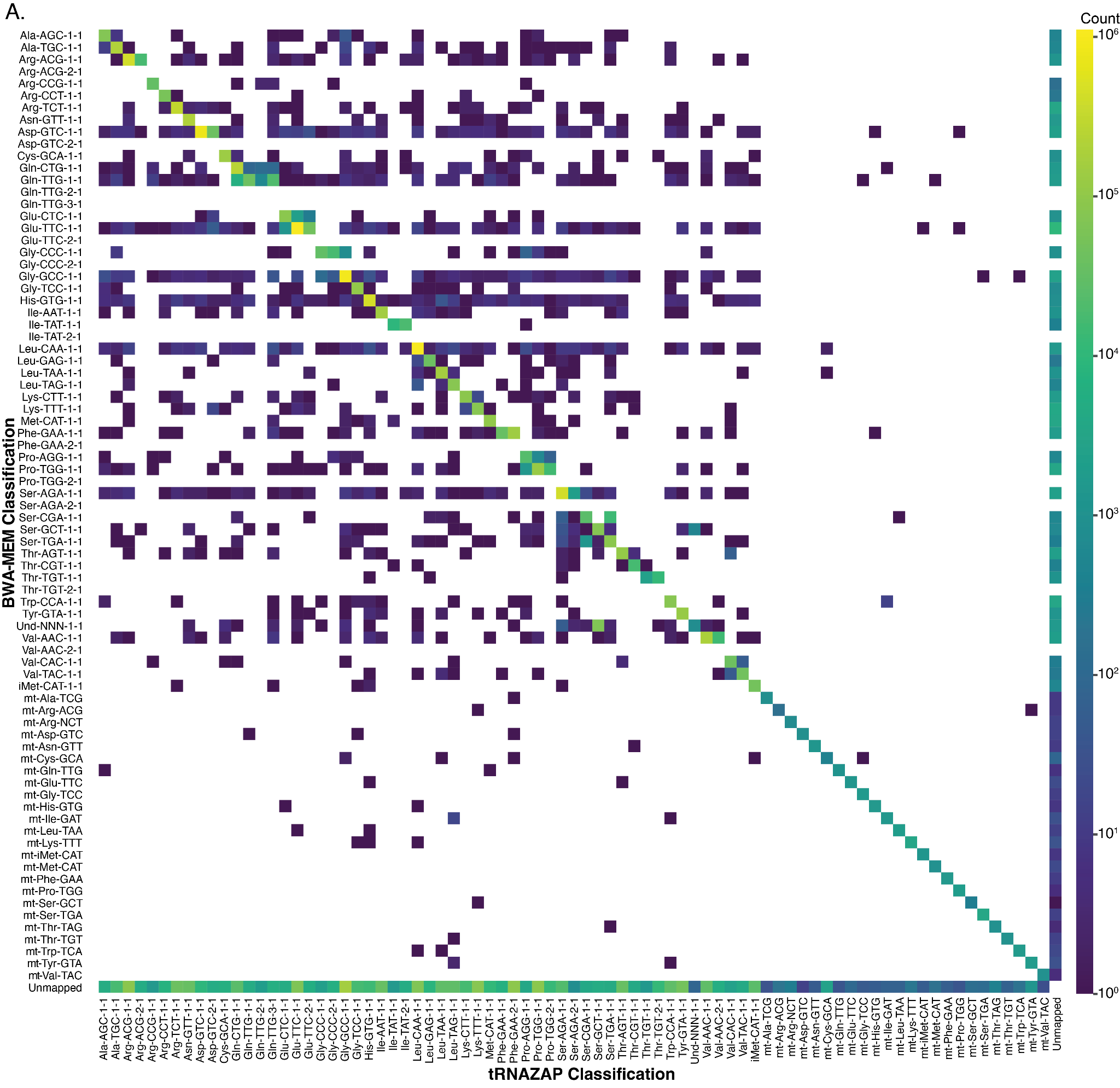

**Supplementary Figure 7. Heatmap showing the aligned read counts for tRNAZAP and BWA-MEM across all *S. cerevisiae* isodecoders. The x-axis shows the reference that tRNZAP aligned the read to, while the y-axis shows the reference that BWA-MEM aligned the read to. The color of each cell represents the total number of reads that had that specific combination of tRNAzap reference and BWA-MEM reference.**

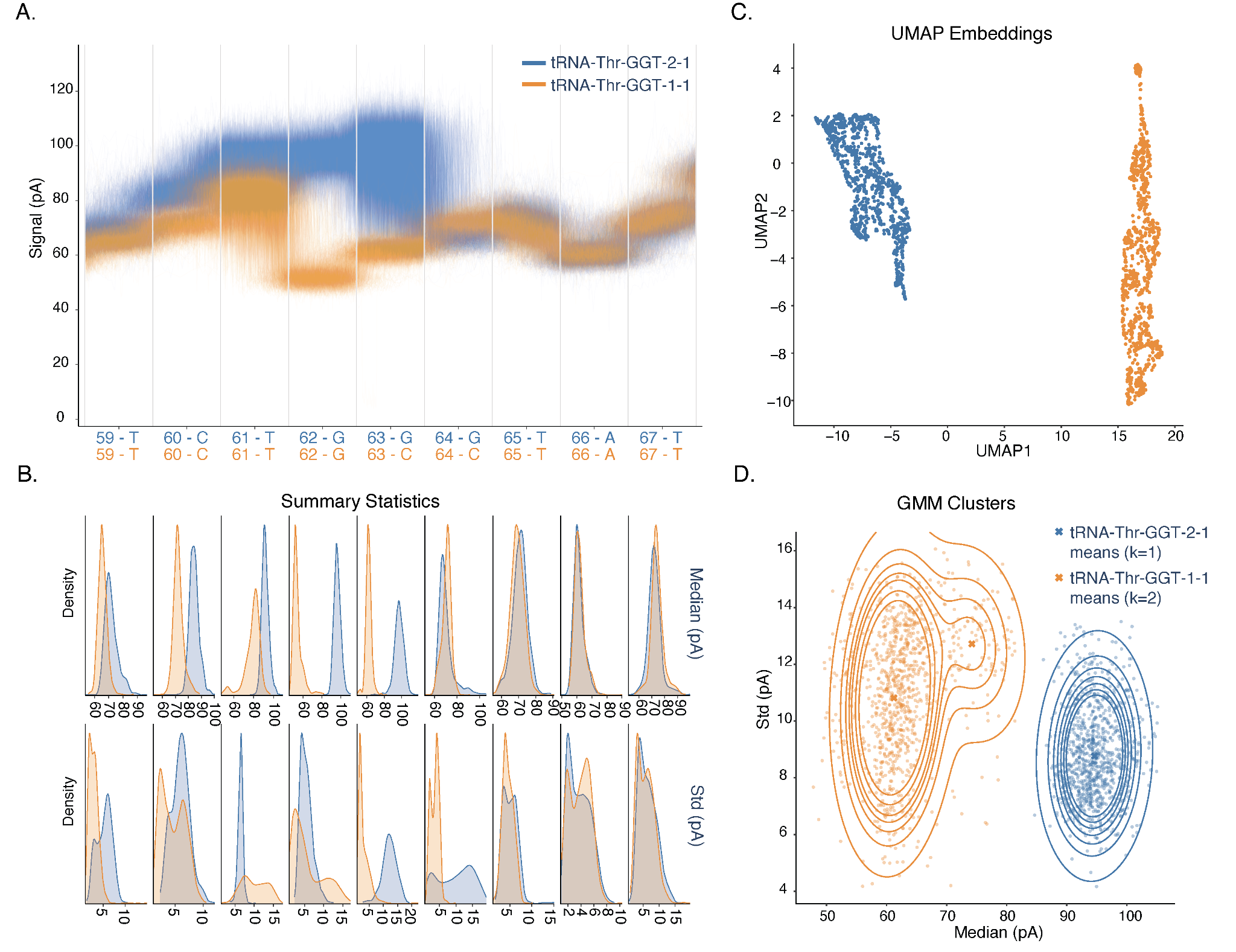

**Supplementary Figure 8. Ionic current comparison between *E. coli* isodecoder variants tRNA-Thr-GGT-1-1 and tRNA-Thr-GGT-2-1.** The two isodecoders differ at positions 63 and 64. A. Raw ionic current traces overlaid across nine positionally matched nucleotide positions centered at position 63, colored by isodecoder. B. Kernel density estimates of per-position median (top) and standard deviation (bottom) of ionic current (pA) for each isodecoder. C. UMAP projection of multi-position ionic current features, revealing population structure between isodecoders. D. Two-dimensional Gaussian mixture models (GMMs) fitted to median and standard deviation features, with cluster means shown for each isodecoder.

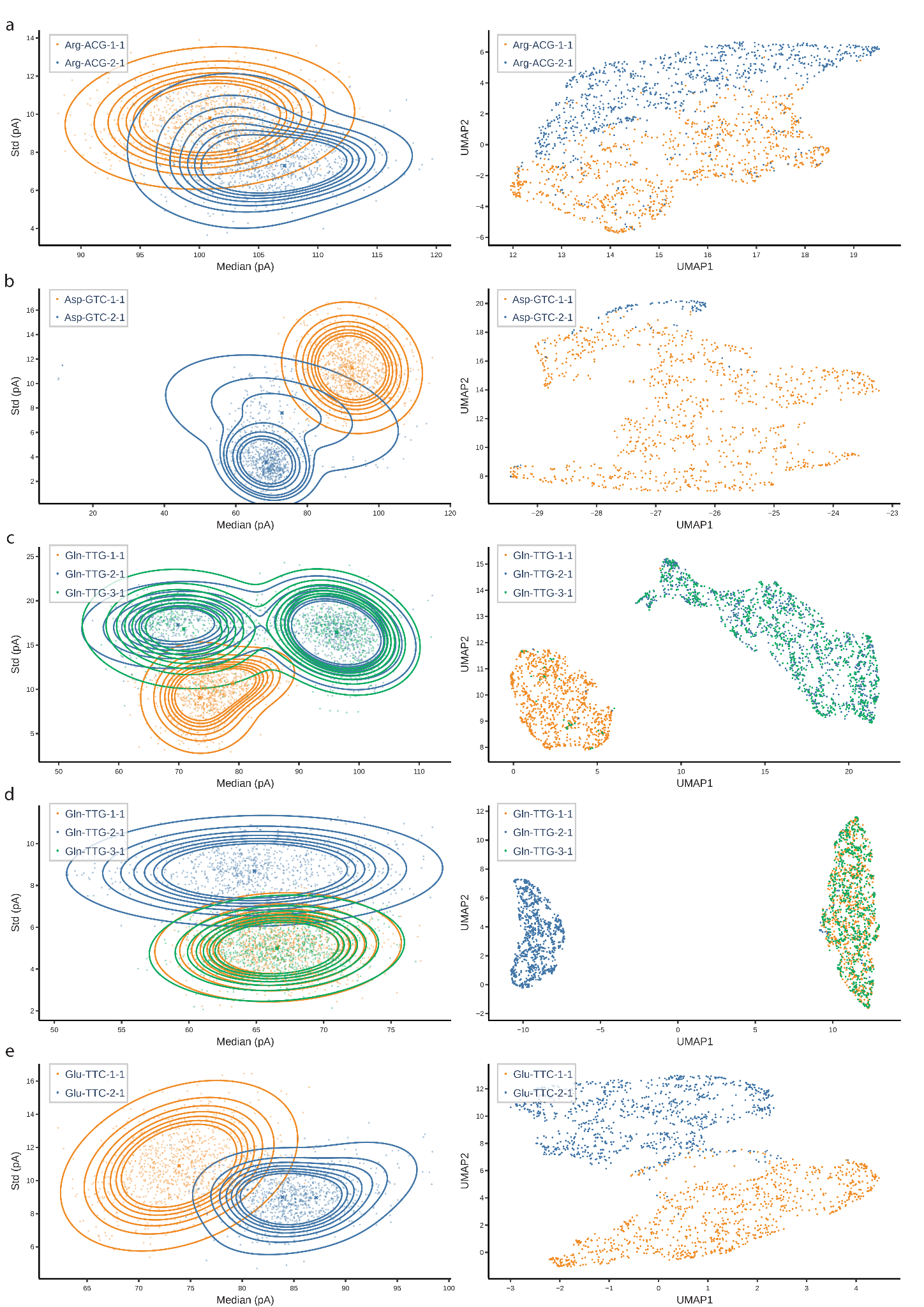

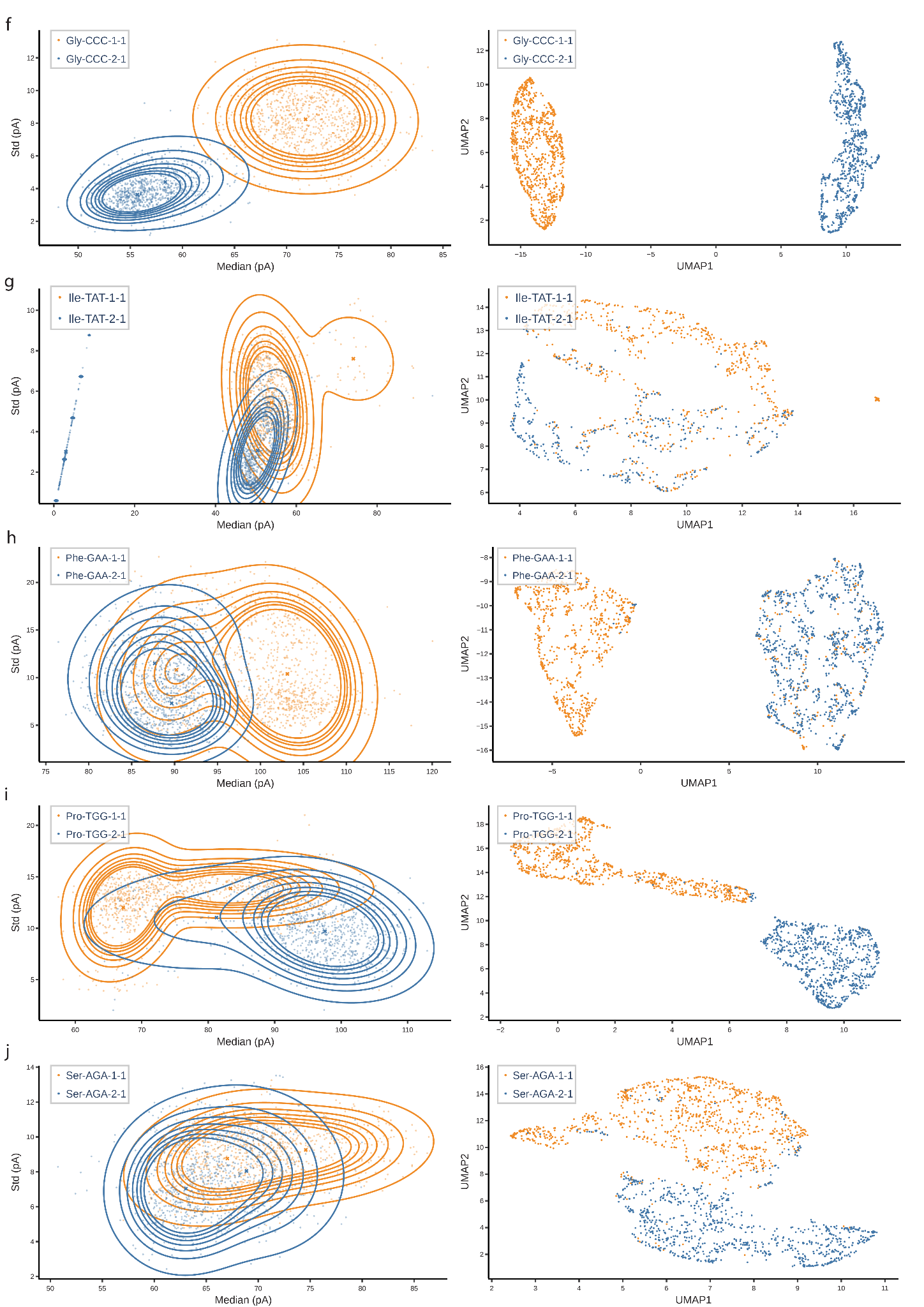

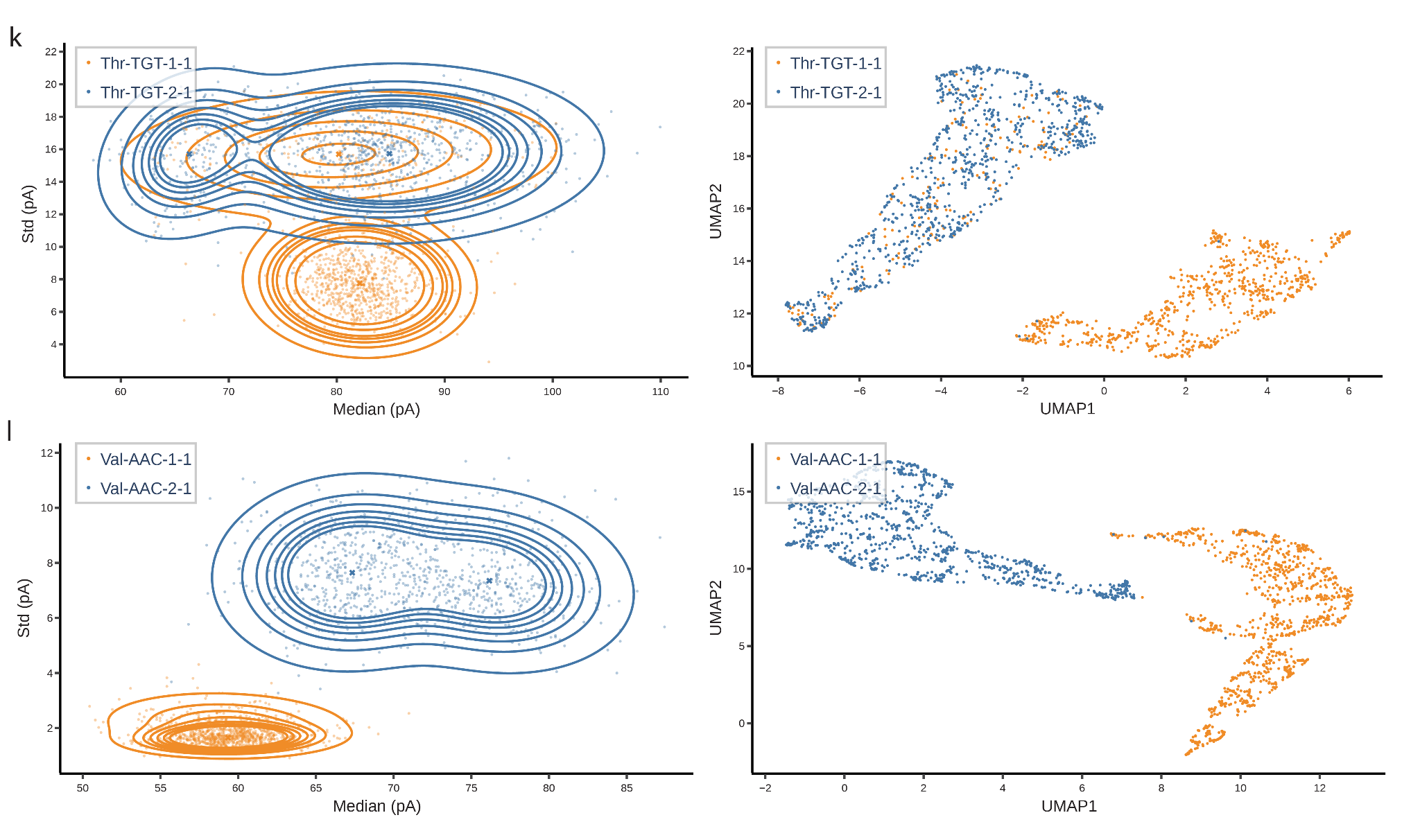

**Supplementary Figure 9.** **Two-dimensional Gaussian mixture models (GMMs) and UMAP of ionic current features for *S. Cerevisiae* isodecoder pairs.** Ionic current signals were extracted from a fixed window centered on the site of sequence difference. A span of consecutive reference positions showing maximal difference between isodecoders was identified and used for feature construction. For GMM fitting, each read was represented as a two-dimensional feature vector comprising the median and standard deviation of the ionic current across this entire span. For UMAP embeddings (left), a higher-dimensional feature vector was constructed per read by computing the median and standard deviation of the ionic current at each individual reference position within the span, then concatenating these values. GMMs with one or two components were fit per isodecoder, with the optimal model selected by the Bayesian Information Criterion (BIC). In the GMM panels, lines denote isomass groups, dots denote individual samples, and crosses denote cluster means.

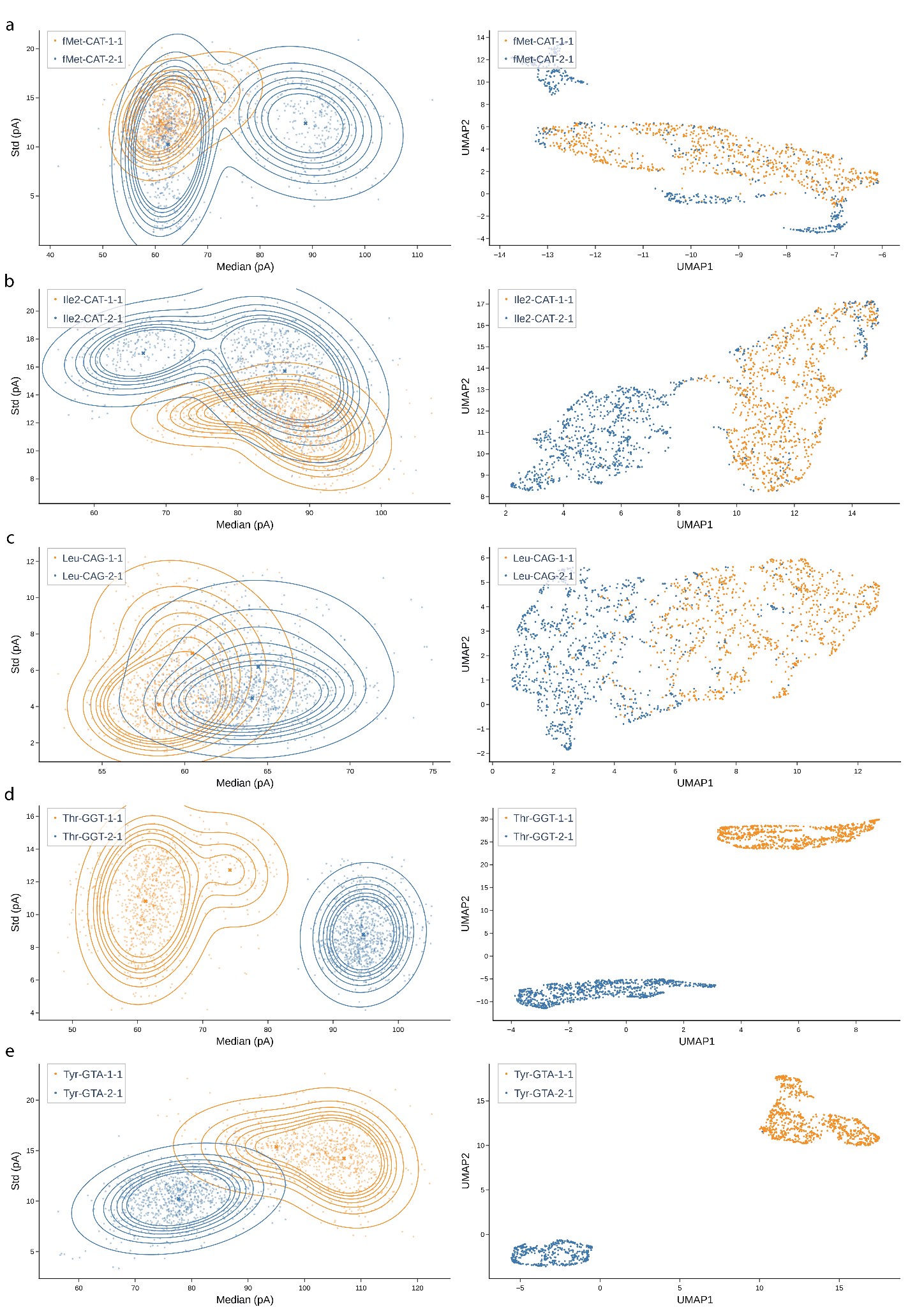

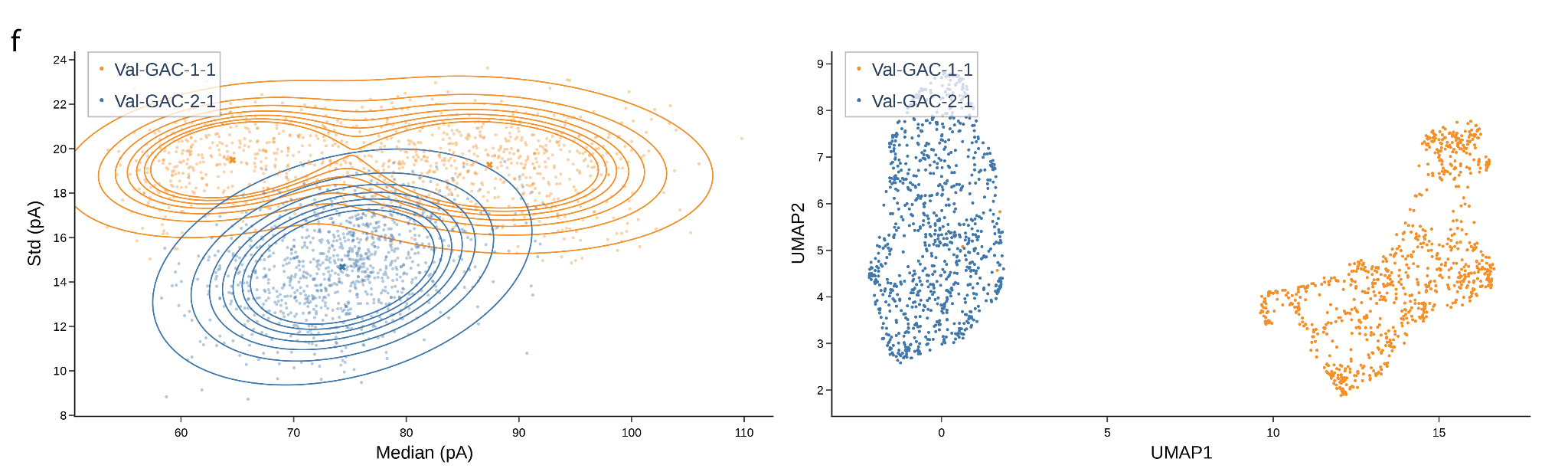

**Supplementary Figure 10.** **Two-dimensional Gaussian mixture models (GMMs) and UMAP of ionic current features for *E. coli* isodecoder pairs.** Ionic current signals were extracted from a fixed window centered on the site of sequence difference. Ionic current signals were extracted from a fixed window centered on the site of sequence difference. A span of consecutive reference positions showing maximal difference between isodecoders was identified and used for feature construction. For GMM fitting, each read was represented as a two-dimensional feature vector comprising the median and standard deviation of the ionic current across this entire span. For UMAP embeddings (left), a higher-dimensional feature vector was constructed per read by computing the median and standard deviation of the ionic current at each individual reference position within the span, then concatenating these values. GMMs with one or two components were fit per isodecoder, with the optimal model selected by the Bayesian Information Criterion (BIC). In the GMM panels, lines denote isomass groups, dots denote individual samples, and crosses denote cluster means.

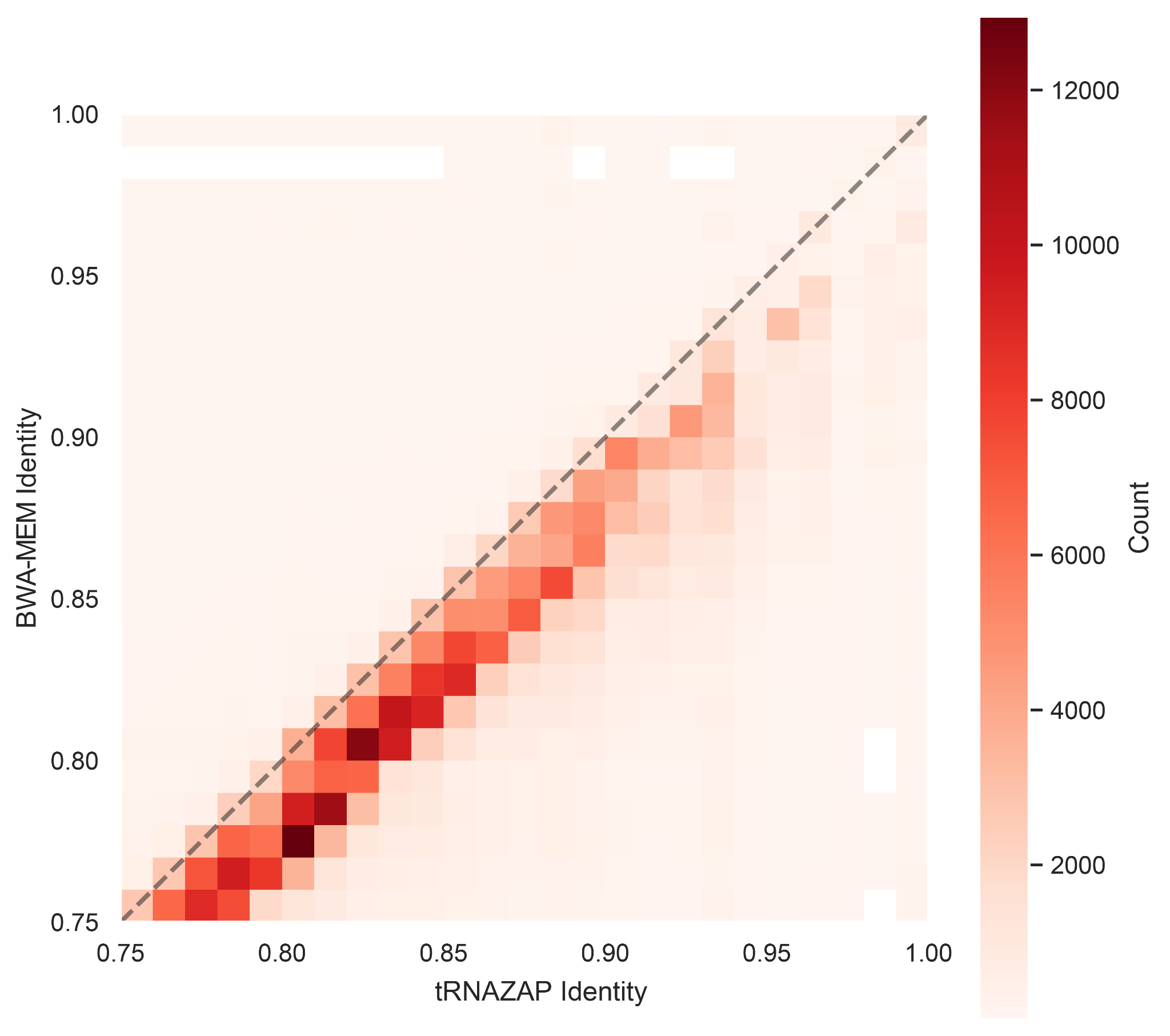

**Supplementary Figure 11.** Comparison of tRNAZAP alignment identity vs. BWA-MEM alignment identity for reads where aligners aligned to different tRNA isodecoders. This comparison is for the *pus*4Δ strain total tRNA sequencing data.

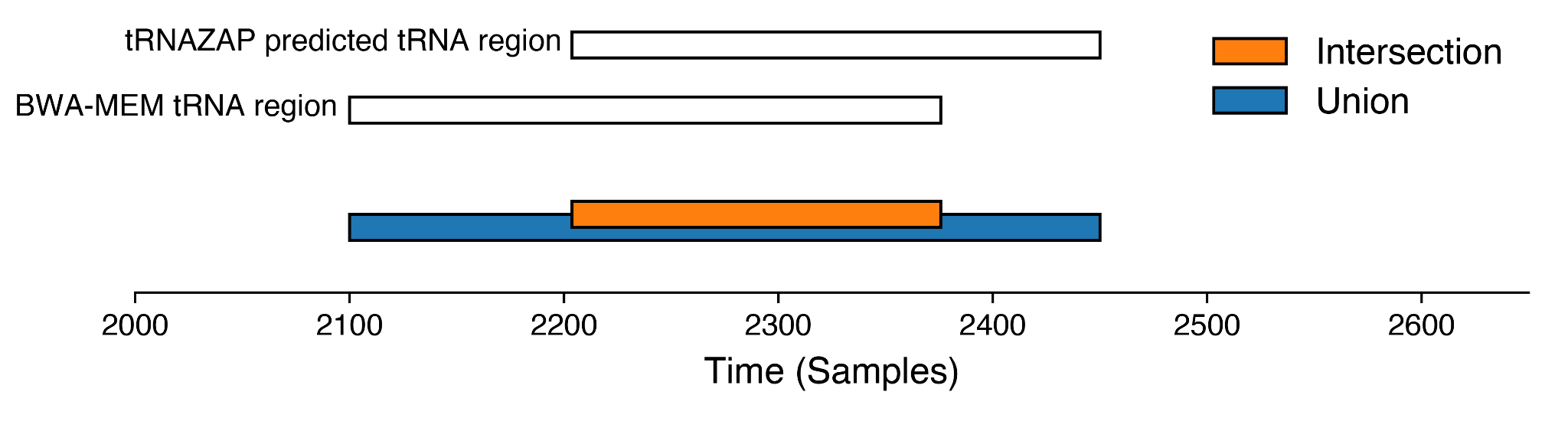

**Supplementary Figure 12.** Intersection-over-Union (IoU) Illustration. The BWA-MEM tRNA region is derived from the BWA-MEM alignment of sequence to the reference and the dorado move table that maps signal points labeled as Time (Samples) here to the sequence. tRNAZAP produces the tRNA segment directly from the ionic current. The overlap (union) of these points measures the agreement while the union (total signal range produced by both tools) provides normalization.

**Supplementary Tables: attached Excel file.**

**Supplementary Table 1. Non-redundant E. Coli / Yeast isodecoders predicted from gTRNAdb**

**Supplementary Table 2. Adapter sequences for IVT and Biological Sequencing**

**Supplementary Table 3. Training / Validation / Test Split per isodecoder**

**Supplementary Table 4. E. Coli Fine Tuning Performance**

**Supplementary Table 5. BWA-MEM and tRNAZAP performance for E. coli isodecoders**

**Supplementary Table 6. E. Coli Phe Enrichment Numbers**

**Supplementary Table 7. BWA-MEM Yeast performance (Identity & Length)**

**Supplementary Table 8. BWA-MEM and tRNAZAP performance for S. cerevisiae isodecoders**

**Supplementary Table 9. JS Divergence**

**Supplementary Table 10. Mitochondrial enrichment identity table**

**Supplementary Table 11. IVT templates for in vitro transcription**

**Supplementary Table 12. Ligation conditions**

**Supplementary Table 13. Yeast and E. Coli IVT Reference for bwa-mem alignment**
